# An open microscopy framework suited for tracking dCas9 in live bacteria

**DOI:** 10.1101/437137

**Authors:** Koen J. A. Martens, Sam P. B. van Beljouw, Simon van der Els, Sander Baas, Jochem N. A. Vink, Stan J. J. Brouns, Peter van Baarlen, Michiel Kleerebezem, Johannes Hohlbein

**Author notes:** These authors contributed equally (Koen J. A. Martens, Sam P. B. van Beljouw).

## Abstract

Super-resolution microscopy is frequently employed in the life sciences, but the number of freely accessible and affordable microscopy frameworks, especially for single particle tracking photo-activation localization microscopy (sptPALM), remains limited. To this end, we designed the miCube: a versatile super-resolution capable fluorescence microscope, which combines high spatiotemporal resolution, good adaptability, low price, and easy installation. By providing all details, we hope to enable interested researchers to build an identical or derivative instrument. The capabilities of the miCube are assessed with a novel sptPALM assay relying on the heterogeneous expression of target genes. Here, we elucidate mechanistic details of catalytically inactive Cas9 (dead Cas9) in live *Lactococcus lactis*. We show that, lacking specific DNA target sites, the binding and unbinding of dCas9 to DNA can be described using simplified rate constants of *k*_bound→free_ = 30−80 s^−1^ and *k*_free→bound_ = 15−40 s^−1^. Moreover, after providing specific DNA target sites via DNA plasmids, the plasmid-bound dCas9 population size decreases with increasing dCas9 copy number via a mono-exponential decay, indicative of simple disassociation kinetics.

---

Fluorescence microscopy has been an integral asset for research in the life sciences for decades and the advent of super-resolution microscopy has been providing additional means to gain insights in biology^1^. In addition to sub-diffractive imaging of cellular structures, super-resolution microscopy can be used to observe the dynamics and interactions of individual proteins in living cells via single particle tracking photo-activatable localization microscopy (sptPALM)^2–4^. Compared to imaging applications, *in vivo* sptPALM has additional experimental challenges to overcome: increased cellular background fluorescence, limited photon budget of single genetically encoded fluorescent proteins and a high time resolution (millisecond range) required to obtain molecular tracks. As full commercial solutions suitable for super-resolution microscopy are costly and restrict the user in their choice of hardware and software, a multitude of simplified custom-builds have emerged^5–11^. We believe, however, that the optimum between the spatiotemporal demands of sptPALM, adaptability, price, and ease of installation has not yet been realized.

To address this need, we designed the miCube: an open-source, simple, modular and versatile microscope suitable for a variety of fluorescence imaging modalities (Fig 1a, Materials and Methods, Supplementary Figure 1, and Supplementary Note 1, https://HohlbeinLab.github.io/miCube). The miCube combines off-the-shelf parts with a custom-designed aluminium body and 3D printed components allowing for sptPALM, sub-diffractive imaging as demonstrated earlier^12^, and total internal reflection fluorescence (TIRF) microscopy. As we minimised the number of necessary components and used pre-aligned arrangements, the miCube does not require extensive expertise in optics or engineering to set up. A detailed description of all components and design choices can be found in Supplementary Note 1.

To demonstrate the capabilities of the miCube and to expand on previous sptPALM studies of CRISPR-Cas^13, 14^, we developed a novel assay that uses a heterogeneous expression system to explore the dynamic nature of DNA-protein interactions in live bacteria and their dependency on protein copy numbers. To this end, we expressed catalytically inactive Cas9 (dead Cas9 or dCas9)^15^ fused to the photo-activatable fluorophore PAmCherry2^16^ under control of the heterogeneous inducible *nisA* promotor^17^ (pLAB-dCas9, Materials and Methods) in *Lactococcus lactis*. In this strain we introduced additional plasmids without (pNonTarget) or with (pTarget) dCas9 DNA target sites (Materials and Methods). Representative cells from each strain showed qualitative differences in the distribution of diffusion coefficients belonging to dCas9 (Fig. 1b).

We analysed the apparent diffusion coefficients as a function of apparent cellular dCas9 copy numbers (Supplementary Note 2). The data of cells containing pNonTarget was fitted with three populations by means of a global fit amongst the different segmented populations (Fig. 1c, Supplementary Figure 2). We attributed the population with the highest apparent diffusion coefficient to freely diffusing dCas9 (*D*_free_* = 1.31 ± 0.05 µm^2^/s). This value is consistent with simulated cytoplasmic diffusion of dCas9-PAmCherry2 (Supplementary Note 3). We found a second population representing a fraction of dCas9 with an apparent diffusion coefficient (*D*_bound_* = 0.13 ± 0.01 µm^2^/s) that is fully determined by the localisation uncertainty of 35-40 nm (Supplementary Note 3), suggesting that even in absence of targets some dCas9 is immobile for longer than ~50 ms, possibly representing partial DNA-sgRNA matching (Supplementary Note 4). In fact, we saw a decrease of the *D*_bound_* population with increasing cellular dCas9 copy numbers from 13 ± 1.5 % to 7.7 ± 1.3%, suggesting that these partial matching sites become increasingly inaccessible to unbound dCas9 (Supplementary Figure 2). We attributed the third population (*D*_trans_ * = 0.46 ± 0.02 µm^2^/s) to transient interactions of dCas9 such as PAM-probing and partial DNA matching that occur on time scales (~10-30 ms) comparable to our acquisition frame time of 10 ms. These time scales are in agreement with previous calculations of transient binding times of dCas9 in *E. coli*^13^. This population of transient interactions is likely to represent temporal averaging of the mobile and various bound states. Assuming a simplified two state model, we determined rate constants of *k*_bound→free_ = 30–80 s^−1^ and *k*_free→bound_ = 15–40 s^−1^ (Supplementary Note 3, Supplementary Figure 3).

The apparent diffusion coefficient histograms of cells containing pTarget were fitted with a combination of the diffusion populations determined from pNonTarget cells for a given range of dCas9 copy numbers and with a new population corresponding to plasmid-bound dCas9 (Fig. 1d; *D*_plasmid_* = 0.17 ± 0.01 µm^2^/s = *D*_bound_* + 0.04 µm^2^/s, which agrees with the expected diffusion coefficient from plasmids of similar size^18^). This *D*_plasmid_* population decreases exponentially (*R*^2^ = 0.98) with increasing apparent cellular dCas9 copy numbers from 20 ± 2.4% at 78 (14–143) copies to 5.3 ± 1.6 % at 642 (571–714) copies (Fig. 1d inset).

In conclusion, we have designed the super-resolution microscope miCube and evaluated its capabilities by performing sptPALM of dCas9 in absence and presence of dCas9 DNA target sites. We showed that the diffusional equilibrium of dCas9 is dynamic and strongly depends on the copy number of dCas9. Our findings further suggest that utilising heterogeneity of an expression system provides an elegant way to obtain insights in any protein-DNA or protein-protein interaction. The high spatiotemporal resolution of the experimental data allowed us to develop a quantitative model of the diffusional equilibrium, proving that the miCube framework is an excellent option even for highly demanding single-molecule super-resolution studies.

**Figure 1:**
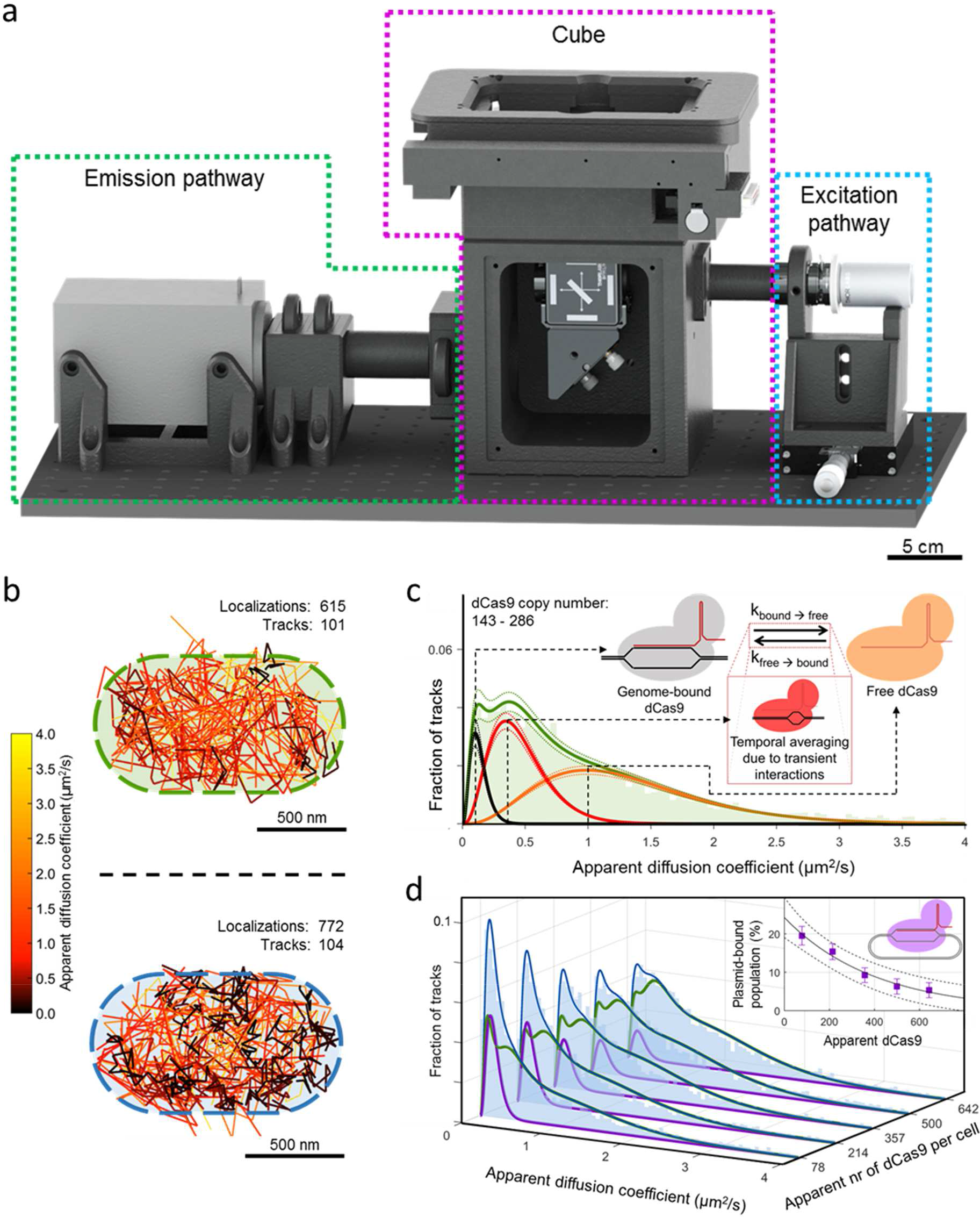
An open-source microscopy framework for sptPALM of dCas9 in *Lactococcus lactis*. (**a**) Render of the open-source miCube super-resolution microscope. The excitation components, main cube, and emission components are indicated in blue, magenta, and green, respectively. A full description of the microscope and all components, including technical drawings, can be found in the Supplementary Note 1. (**b**) Single particle tracking of dCas9 on the miCube in individual *L lactis* cells. The tracks are colour-coded to reflect the apparent diffusion coefficient. Top: a pNonTarget-containing cell. Bottom: a pTarget-containing cell. (**c**) Global fitting of the apparent diffusion coefficients reveals three distinct populations in pNonTarget-containing cells. The apparent diffusion coefficient histogram for the 143–286 dCas9 copy number range is shown in green (bin width = 0.05 µm^2^/s). The average population sizes of the three populations are indicated with a solid line, and the corresponding 95% confidence interval of the fit (least-squares) of the populations is shown in dotted lines. The populations are attributed to genome-bound dCas9 (grey), transient interactions that lead to temporal averaging (red), and freely diffusing dCas9 (orange). Interaction between the three states is shown in the figure, where k_bound→free_ = 30–80 s^−1^ and k_free→bound_ = 15–40 s^−1^, determined from all dCas9 copy number ranges. (**d**) In pTarget-containing *L. lactis* cells, the apparent diffusion coefficient histograms (bin width = 0.05 µm^2^/s) are fit with the appropriate combined population of the pNonTarget-containing cells (green), and with a population corresponding to target-bound dCas9 (purple). The population size of the plasmid-bound dCas9 decreases exponentially as a function of the cellular dCas9 copy number (inset). The error bar is based on the 95% confidence interval of the histogram fits (least-squares); the dotted lines of the exponential decrease is based on the 95% confidence interval of the equation fit (least-squares). Data shown in (c) and (d) comprises of four individual repeats measured over three days. For pNonTarget, we analysed 628 cells leading to 224.050 localisations forming 32.971 tracks of at least 4 localisations. For pTarget, we analysed 428 cells, 220.635 localisations, and 31.439 tracks.

## Materials and Methods

### Biological methods

#### Strain preparation and plasmid construction

*Lactococcus lactis* NZ9000 was used throughout the study. NZ9000 is a derivative of *L. lactis* MG1363^1^ in which the chromosomal *pepN* gene is replaced by the *nisRK* genes that allow the use of the nisin-controlled gene expression system^2^. Cells were grown at 30°C in GM17 medium (M17 medium (Tritium, Eindhoven, The Netherlands) supplemented with 0.5% (w/v) glucose (Tritium, Eindhoven, The Netherlands) without agitation.

#### DNA manipulation and transformation

Vectors used in this study are listed in Supplementary Table 1. Oligonucleotides (Supplementary Table 2) and primers (Supplementary Table 3) were synthesised at Sigma-Aldrich (Zwijndrecht, The Netherlands). Plasmid DNA was isolated and purified using GeneJET Plasmid Prep Kits (Thermo Fisher Scientific, Waltham, MA, USA). Plasmid digestion and ligation were performed with Fast Digest enzymes and T4 ligase respectively, according to the manufacturer’s protocol (Thermo Fisher Scientific, Waltham, MA, USA). DNA fragments were purified from agarose gel using the Wizard SV gel and PCR Clean-Up System (Promega, Leiden, The Netherlands). Electro competent *L. lactis* NZ9000 cells were generated using a previously described method^3^. Prior to electro-transformation, ligation mixtures were desalted for one hour by drop dialysis on a 0.025 µm VSWP filter (Merck-Millipore, Billerica, US) floating on MQ water. Electro-transformation was performed with GenePulser XcellTM (Bio-Rad Laboratories, Richmond, California, USA) at 2 kV and 25 µF for 5 ms. Transformants were recovered for 75 minutes in GM17 medium supplemented with 200 mM MgCl_2_ and 2 mM CaCl_2_. Chemically competent E. coli TOP10 (Invitrogen, Breda, The Netherlands) were used for transformation and amplification of the Pnis-dCas9-PAmCherry2-containing pUC16 plasmid (Supplementary Figure 4). Antibiotics were supplemented on agar plates to facilitate plasmid selection: 10 µg/ml chloramphenicol (for pTarget/pNonTarget) and 10 µg/ml erythromycin (for pLAB-dCas9). Screening for positive transformants was performed using colony PCR with KOD Hot Start Mastermix according to the manufacturer’s instructions (Merck Millipore, Amsterdam, the Netherlands). Electrophoresis gels were made with 1% agarose (Eurogentec, Seraing, Belgium) in tris-acetate-EDTA (TAE) buffer (Invitrogen, Breda, The Netherlands). Plasmid digestions were compared with *in silico* plasmid digestions (Benchling; https://benchling.com).

#### pLAB-dCas9 plasmid construction

The sequence of Pnis-dCas9-PAmCherry2 flanked by XbaI/SalI restriction sites (Supplementary Figure 4, Supplementary Note 5) was synthesized by Baseclear (Baseclear B.V., Leiden, The Netherlands), and cloned in a pUC16 plasmid. After transformation in *E. coli*, the plasmid was isolated and digested with XbaI and SalI to obtain the Pnis-dCas9-PAmCherry2 fragment. The Cas9 expression module was removed from the pLABTarget expression vector^4^ by digestion with XbaI and SalI and replaced by the XbaI-SalI fragment containing Pnis-dCas9-PAmCherry2. The single-stranded guide RNA (sgRNA) for targeting *pepN* was constructed and inserted according to earlier described protocol^4^ to yield the pLAB-dCas9 vector (Supplementary Figure 5). The plasmids used in this study, and vector maps for pLABTarget and pLAB-dCas9 are available upon request.

#### pTarget and pNonTarget plasmid construction

The plasmid with binding sites for dCas9 (pTarget) was established by engineering five *pepN* target sites in the pNZ123 plasmid^5^. To this end, two single-stranded oligonucleotides (10 µl of 100 µM, each) that upon hybridization form the target sequence of the *pepN*-targeting sgRNA were incubated in 80 µl annealing buffer (10 mM Tris [pH = 8.0] and 50 mM NaCl) for 5 minutes at 100°C, followed by gradual cooling to room temperature. The annealed multiplexed oligonucleotide was cloned in HindIII-digested pNZ123, yielding a derivative that contains five *pepN* target sites. HindIII re-digestion was prevented by flanking the *pepN* DNA product by different base pairs, changing the HindIII site. Plasmids with five *pepN* target sites were designated pTarget (Supplementary Figure 7). Plasmids without the pepN target sites (the original pNZ123 plasmids) were designated pNonTarget. The vector maps for pTarget and pNonTarget are available upon request.

#### Construction of strains harbouring both pLAB-dCas9 and pTarget/pNonTarget

Electro competent *L. lactis* NZ9000 cells^3^ harbouring pLAB-dCas9 were transformed with pTarget or with pNonTarget and subsequently used for sptPALM or stocked at −80°C.

### Single molecule microscopy

#### Sample preparation

Cells were grown overnight from cryostocks in chemically defined medium for prolonged cultivation (CDMPC)^6^. The overnight cultures were sub-cultured (1:50 (v/v)) in CDMPC until exponential phase was reached (after 4 hours of growth) at 30°C. Cultures were then incubated for 1.5h at 30°C with 0.4 ng/mL nisin (Sigma-Aldrich, Zwijndrecht, The Netherlands) to induce dCas9 expression. 0.5 µg/mL ciprofloxacin (Sigma-Aldrich, Zwijndrecht, The Netherlands) was added to inhibit further cell division and DNA replication^7^. Then, cells were centrifuged (3500 RPM for 5 minutes; SW centrifuge (Froilabo, Mayzieu, France) with a CENSW12000024 swing-out rotor fitted with CENSW12000006 15 ml culture tube adaptors) and washed two times by gentle resuspension in 5 mL phosphate-buffered saline (PBS; Sigma-Aldrich, Zwijndrecht, The Netherlands). After removal of the supernatant, cells were resuspended in ~10-50 µL PBS from which 1-2 µL was immobilized on 1.5% 0.2 µm-filtered agarose pads between two heat-treated glass coverslips (Paul Marienfeld GmbH & Co. KG, Lauda-Königshofen, Germany; #1.5H, 170 µm thickness). Heat treatment of glass coverslips involves heating the coverslips to 500°C for 20 minutes in a muffle furnace to remove organic impurities.

#### Experimental settings

All imaging was performed on the miCube as described further in Supplementary Note 1. A 561 nm laser with ~0.12 W/cm^2^ power output was used for HiLo-to-TIRF illumination with 4 ms stroboscopic illumination^8^ in the middle of 10 ms frames. Low-power UV illumination (µW/cm^2^ range) was used and increased during experiments to ensure a low and steady number of fluorophores in the sample until exhaustion of the fluorophores. A UV-increment scheme was consistently used for all experiments (Supplementary Table 4). No emission filter was used except for the TIRF filter (Chroma ZET405/488/561m-TRF). The raw data was acquired using the open source Micro-Manager software^9^. During acquisition, 2×2 binning was used, which resulted in a pixel size of 128×128 nm. The camera image was cropped to the central 512×512 pixels (65.64 × 65.64 µm). For sptPALM experiments, frames 500-55,000 were used for analysis, corresponding to 5-550 seconds. This prevented attempted localization of overlapping fluorophores at the beginning, and ensured a set end-time. 200-300 brightfield images were recorded by illuminating the sample at the same position as during the measurement. For the brightfield recording, we used a commercial LED light (INREDA, IKEA, Sweden) and a home-made diffuser from baking paper.

#### Localization

To extract single molecule localizations, a 50-frame temporal median filter (https://github.com/marcelocordeiro/medianfilter-imagej) was used to correct background intensity from the movies^10^. In short, the temporal median filter determines the median pixel value over a sliding-window of 50 pixels to determine the median background intensity value for a pixel at a specific position and time point. This value is subtracted from the original data, and any negative values are set to 0. In the process, all pixels are scaled according to the mean intensity of each frame to account for shifts in overall intensity. The first and last 25 frames from every batch of 8096 frames are removed in this process.

Single particle localization was performed via the ImageJ^11^/FIJI^12^ plugin ThunderSTORM^13^ with added phasor-based single molecule localization algorithm (pSMLM^14^). Image filtering was done via a difference-of-Gaussians filter with Sigma1 = 2 px and Sigma2 = 8 px. The approximate localization of molecules was determined via a local maximum with a peak intensity threshold of 8, and 8-neighbourhood connectivity. Sub-pixel localization was done via phasor fitting^14^ with a fit radius of 3 pixels (region-of-interest of 7-by-7 pixels). Custom-written MATLAB (The MathWorks, Natick, MA, USA) scripts were used to combine the output files from ThunderSTORM.

#### Cell segmentation

A cell-based segmentation on the localization positions was performed. First, a watershed was performed on the average of 300 brightfield-recorded frames of the cells. The watershed was done via the Interactive Watershed ImageJ plugin (http://imagej.net/Interactive_Watershed). Second, the localizations were filtered whether or not they fall in a pixel-accurate cell outline. If they do, a cell ID is added to the localization information.

#### Tracking and fitting of apparent diffusion coefficient histogram

A tracking procedure was performed in MATLAB, using a modified Particle Point Analysis script^15^ (https://nl.mathworks.com/matlabcentral/fileexchange/42573-particle-point-analysis) with a tracking window of 8 pixels (1.0 µm) and memory of 1 frame. Localizations corresponding to different cells were excluded from being part of the same track. As the tracking window is of similar size as the cells itself, in practice all localizations in a cell are linked together in a track if they appear in successive frames, or skip maximum one frame.

An apparent diffusion coefficient, *D*\*, was then calculated for each track from the mean-squared displacement (MSD) of single-step intervals^16^. In short, for every track with at least 4 localizations, the *D*\* was calculated by calculating the mean square displacement between the first four steps (taking skipped steps due to a memory of 1 frame into account in the calculated distance), and taking the average of that. These *D*\* values were globally fitted with a custom-written MATLAB script, which iteratively fits (a combination of non-linear regression and non-linear least-squares) a combination of 3 populations with a variable *D*\*-value and population size on multiple datasets. In short, equation (1) was globally fitted to multiple histograms of the datasets.

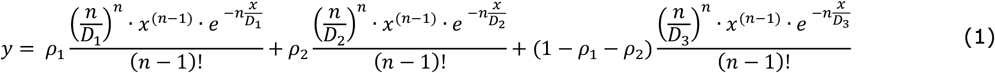

Where *ρ*_1_ and *ρ*_2_ represent relative occupancies of the three populations; *D*_1_, *D*_2_, and *D*_3_ the *D*\*-values, *n* the number of steps in the trajectory (set to four in this study), *y* the intensity of the histogram, and *x* the *D*\*-value of the histogram. *D*_1_, *D*_2_, and *D*_3_ were kept constant in the global fit; *ρ* and *ρ*_2_ were allowed to vary between datasets. A 95% confidence interval on all parameters was acquired by analysing the goodness-of-fit of the curve to the datasets.

#### miCube drift quantification

We characterised the positional stability of the miCube via super-resolution measurements of GATTA-PAINT 80R DNA-PAINT nanorulers (GATTAquant GmbH, Germany). We imaged the nanorulers in total internal reflection (TIR) mode using a 561 nm laser (~7 mW) with a frame time of 50 ms using 2×2 pixel binning on the Andor Zyla 4.2 PLUS sCMOS. Astigmatism was enabled by placing a 1000 mm focal length astigmatic lens (Thorlabs) 51 mm away from the camera chip. A video of 10.000 frames was recorded via the MicroManager software^9^.

After recording the movie, we first localized the *x, y*, and *z*-positions of the point spread functions of excited DNA-PAINT nanoruler fluorophores with the ThunderSTORM software^13^ for ImageJ^11^ with the phasor-based single molecule localization (pSMLM) add-on^14^. The ThunderSTORM software was used with the standard settings, and a 7 by 7 pixel region of interest around the approximate centre of the point spread functions was used for pSMLM. To determine the *z*-position, we compared the astigmatism of the point-spread function to a pre-recorded calibration curve recorded using immobilized fluorescent latex beads (560 nm emission peak, 50 nm diameter).

After data analysis we performed drift-correction in the lateral plane using the cross-correlation method of the ThunderSTORM software. The cross-correlation images were calculated using 10x magnified super-resolution images from a sub-stack of 100 original frames. The fit of the cross-correlation was used as drift of the lateral plane. Drift of the axial plane was analysed by taking the average *z*-position of all fluorophores, assuming that all DNA-PAINT nanorulers are fixed to the bottom of the glass slide.

## Acknowledgments

K.J.A.M. is funded by a VLAG PhD-fellowship grant awarded to J.H. J.H. acknowledges funding from the Innovation Program Microbiology Wageningen (IPM-3). S.v.d.E is funded by the BE-Basic R&D program, which was granted a FES subsidy from the Dutch Ministry of Economic affairs.

## Author contributions

K.J.A.M., S.B., and J.H. designed, built and characterised the miCube setup. K.J.A.M. and S.P.B.v.B. recorded and analysed the experimental single molecule data. J.H., S.v.d.E and P.v.B. envisioned using *L. lactis*, dCas9, fluorescent proteins and (Non-)Target cells to conduct super-resolution single molecule studies. S.P.B.v.B, S.v.d.E., P.v.B., and M.K. designed the DNA vectors used in this study. S.P.B.v.B. and S.v.d.E. assembled the DNA vectors and transformed cells. J.N.A.V and K.J.A.M. wrote additional software for data analysis. J.N.A.V and S.J.J.B. provided reagents and expertise for setting up the single molecule assays. K.J.A.M., S.P.B.v.B., and J.H. wrote the manuscript with input from all authors. All authors contributed to the design of the assays. J.H. initialised the study and the collaborations, and supervised all aspects of the study.

## Supplementary information

### Supplementary information Supplementary Note 1: Detailed description miCube

Custom-build instruments or partial instruments^17^ have been presented before, such as the adaptation of commercial microscopes to add functionalities^18^, an open-frame microscope which provide dual-emission super-resolution^19^, simplified and low-cost super-resolution microscopes^20–24^, and a lab-on-a-chip design using polydimethylsiloxane (PDMS) rather than glass-based optics^25^. We designed the miCube to be easy to set up and use, while retaining a high level of versatility. The instrument and its design choices will be described in three parts: the excitation path; the emission path, and the ‘cube’ connecting the sample with the excitation and emission paths. Throughout this description, we will refer to numbered parts as shown in Supplementary Figure 1a and c and described in Supplementary Table 5. The information on the miCube presented here can also be found on https://HohlbeinLab.github.io/miCube/component_table.html. The instrument is fully functional in ambient light, due to a fully enclosed sample chamber, illumination pathway and emission pathway. Moreover, the miCube has a small footprint: the final design of the miCube, excluding the lasers and controllers, fits on a 300 × 600 mm Thorlabs breadboard. We placed the whole ensemble in a transparent polycarbonate box (MayTec Benelux, Doetinchem, The Netherlands) to minimize airflow disturbing the setup during experiments.

#### A. Excitation path

The excitation path is designed to be both robust and easy to align and adjust. The four laser sources located in an Omicron laser box are combined and guided via a single mode fibre towards a reflective collimator (nr. 18) ensuring a well-collimated beam. The reflective collimator is attached directly to an aperture (nr. 17), a focusing lens (nr. 16, 200 mm focus length), and an empty spacer (nr. 12). This excitation ensemble is placed in the 3D-printed piece specifically designed to hold the assembly into place (nr. 13). This holder is then attached to a right-angled mounting plate (nr. 14), which is placed on a 25mm translation stage (nr. 15). The translation stage should be placed at such a position on the breadboard that the focusing lens (nr. 16) is exactly 200 mm separated from the back-focal plane of the objective when following the laser path.

Easy alignment and adjustment are ensured by isolating the three axes of movement of this excitation ensemble (Supplementary Figure 6). Adjustments of distance from objective is achieved by moving the collimator ensemble (nrs. 12, 16-18) inside its holder (nr. 13). Height of the path can be adjusted via a bracket clamp that supports the collimator ensemble (nrs. 13 and 14), and the horizontal alignment can be adjusted via a translation stage where the bracket clamp rests on (nr. 15).

Additionally, the translation stage (nr. 15) can be used to enable highly inclined illumination (HiLo) or total internal reflection (TIR). The stage allows fine and repeatable adjustment of the excitation beam position on the back focal plane of the objective. By aligning the excitation beam in the centre of the objective, the microscope will act as a standard epifluorescence instrument. If the excitation beam is aligned towards the edge of the back focal plane, the miCube will operate in HiLo or TIR.

#### B. Cube and sample mount

The central component of the miCube is the cube (nr. 5) that connects excitation path, emission path, and the sample. The cube is manufactured out of a solid aluminium block maximising stability and minimising effects of drift due to thermal expansion (Supplementary Note 6). Black anodization of the block prevents stray light and unwanted reflections. The illumination path is further protected from ambient light by screwing a 3D-printed cover (nr. 11) on the side of the cube, as well as a door to close the cube off.

Next, the ‘dichroic mirror - full mirror’ part is assembled (nrs. 6-10). The dichroic mirror unit (nr. 7) consists of a dichroic mount that is magnetically attached to an outer holder. On the side of the dichroic mirror unit, opposing the excitation path, a neutral density filter (nr. 6) is placed to prevent scattered non-reflected light entering the miCube thereby minimizing background signal being recorded by the camera. At the bottom of the dichroic mount assembly, a TIRF filter (nr. 8) is placed to remove scattered back-reflected laser light from entering the emission pathway. This ensembled dichroic mirror unit is screwed via a coupling element (nr. 9) to a mirror holder containing a mirror placed at a 45° angle (nr. 10), which reflects the emission light from the objective to the camera. This completed ‘dichroic mirror - full mirror’ part is screwed into the backside of the miCube via two M6 screws, which hold the ensemble into place and directly in line with the excitation path (nrs. 12-18), the objective (nr. 3), and the tube lens (nr. 30).

Then, an objective (nr. 3) (Nikon 100x oil, 1.49 NA, HP, SR) is directly screwed into an appropriate thread on top of the cube. Around the objective, a sample mount (nr. 4) is screwed on top of the cube, which is designed to ensure correct height of the sample with respect to the parfocal distance of the chosen objective. We opted for using a sample mount, as it can be easily swapped for another to retain freedom in peripherals. For example, only the height of the sample mount has to be changed if the objective has a different parfocal distance as the one used here. We designed two different sample mounts (nr. 4a, 4b). The first one can hold an xy-translation stage with z-stage piezo insert (nr. 2), to enable full spatial control of the sample (nr. 4a). The second one only holds the z-stage piezo insert, which decreases instrument cost (nr. 4b). In any case, the xy-translation stage with z-stage piezo insert, or only the z-stage piezo insert is screwed in place into corresponding threaded holes in the sample mount. A glass slide holder (nr. 1) is made from aluminium to fit inside a 96-wells-holder like the *z*-stage (nr. 2).

#### C. Detection path

A tube lens ensemble is made (nrs. 27-30) which houses a 200 mm focal length tube lens (Thorlabs) in a 3D-printed enclosure which provides space to slot in an emission filter (nrs. 27,28). This ensemble is then attached directly to the miCube by screwing it into place with four M6 screws. The alignment of the tube lens is therefore exactly in line with the emission light, as the centre of the full mirror (nr. 10) is at the same height of the tube lens. The direction of the emission light can be aligned, which can simply be achieved by tuning the angle of the full mirror (nr. 10).

A cover (nr. 25) is attached to the tube lens ensemble to ensure darkness of the emission path, which is connected to the tube lens by a 3D-printed connector piece (nr. 26). On the other end of the cover, a 3D-printed holder for 2 astigmatic lenses (nr. 21) is placed and screwed into place in the breadboard. Astigmatic lenses (nrs. 22-24) can optionally be used to enable 3D super-resolution microscopy^26^. They can be easily changed for lenses with a different focal length or with empty holders. With this, astigmatism can be enabled or disabled, and a choice between more accurate z-positional information with a lower total z-range, or less accurate information with a larger range can be made. The Andor Zyla 4.2 PLUS camera (nr. 19) is placed behind the astigmatic lens holder, and is positioned in a 3D-printed camera mount (nr. 20) to ensure correct height and position of the camera, so that the focus of the tube lens is at the camera chip. We chose for a scientific Complementary Metal-Oxide Semiconductor (sCMOS) camera to take advantage of a larger field of view and increased temporal resolution compared to the more traditional electron-multiplying charge coupled device (EMCCD) cameras^27^.

Note that the length of the cover (nr. 25) and the alignment of the holes at the feet of the 3D-printed astigmatic lens holder (nr. 21) are dependent on the focal length of the tube lens, and of the position of the chosen camera chip with regards to the 3D-printed mount for the camera. The pieces used here were designed for the Andor Zyla 4.2 PLUS, a 200 mm focal length tube lens, and the used custom-designed camera mount (nr. 20).

### Supplementary Note 2: Estimating the copy number of dCas9

The total copy number of dCas9 in a cell is not identical to the number of tracks found in each cell. Firstly, the UV illumination (405 nm wavelength) on the miCube required to photo-activate PAmCherry2 is not homogeneous over the complete field of view. To correct for this, a value for the average UV illumination experienced by each *L. lactis* cell is calculated. For this, a map of the UV intensity is made by placing a mirror on top of the objective and measuring the reflected scattering of the UV signal. Then, the mean UV intensity in the pixels corresponding to a cell according to the segmented brightfield images is stored. The cellular apparent dCas9 copy number is corrected for this normalized mean cellular UV intensity. Moreover, the cellular apparent dCas9 copy number was corrected for the average maturation grade of PAmCherry1, which is ~70%^28^. We assume the maturation grades of PAmCherry1 and PAmCherry2 to be similar.

### Supplementary Note 3: Model simulations

The theoretical diffusion coefficient of the dCas9-PAmCherry2 construct is 36 - 43 µm^2^/s in water, using the Stokes-Einstein equation^29^, and assuming a hydrodynamic radius of 5 - 6 nm (PDB 5CZZ for Cas9^30^, and PDB 3KCT^28^ for PAmCherry2). This is 18 - 22 times higher than the found real diffusion coefficient of 1.95 µm^2^/s of dCas9-PAmCherry2. However, the increased viscosity and the molecular crowding in the cytoplasm have to be considered. For a construct this size, it is expected that the D_0_/D_cyto_ ratio is ~20.31

Brownian motion of 33.000 tracks of particles was simulated with diffusion coefficients of 0 µm^2^/s, 0.43 µm^2^/s and 1.95 µm^2^/s in a 8:35:57 ratio, with an added 36.5 nm localization uncertainty, in a modelled cell consisting of 2 half-spheres with radius 0.5 µm connected by a straight cylinder of length 0.5 µm and radius 0.5 µm. 10 ms frames were calculated from 10 sub-frames in which the movement of the particles was Gaussian distributed according to the given diffusion coefficient. Each particle was given at least 100 ms of diffusion prior to measurement to mimic equilibrium that molecules may reach. Identical fitting was performed as in the rest of the study.

Performing these theoretical Brownian motion simulations of the dCas9-PAmCherry2 construct showed that this simulation corresponds to fitted apparent diffusion coefficients of 0.12-0.13 µm^2^/s, 0.44-0.48 µm^2^/s, and 1.26-1.36 µm^2^/s in a 9-10:44-48:51-56 ratio (Supplementary Figure 8). This is identical to the fitted pNonTarget apparent diffusion coefficient histogram (Fig. 1c).

A second set of simulations was performed using only real diffusion coefficients of 0 µm^2^/s and 1.95 µm^2^/s (Supplementary Figure 3). However, this simulation was extended such that rate constants (corresponding to [free to bound] transition and [bound to free] transition) determine switching time between the states. By increasing the rate constants, the resulting histogram of apparent diffusion coefficients becomes more convoluted. This is in line with the hypothesis that the transient interactions are a convolution of the bound population and the freely diffusion population (main text). This simulation can thus be used to approximate the rate constants (Supplementary Figure 3 g-k), although fitting this two-state simulation with the experimental data leads to larger residuals than the three-state simulation described above.

### Supplementary Note 4: Unspecific interactions in the *L. lactis* genome

Interactions of dCas9 with genomic DNA are very complex to quantify, but we do offer some discussion on the matter. It has been reported using an *in vitro* assay that the DNA binding equilibrium of dCas9 shifts quickly to unbound dCas9 with an increasing number of mismatches^32^, but there is still some dCas9 bound to DNA even with minimal sgRNA-DNA complementarity. Because we are observing interactions at the short timescale of ~40 ms, short DNA complementarity will have an effect on the occupancy of this population.

Since we see a dependency of dCas9 copy numbers on the genome-bound population of dCas9 (Supplementary Figure 2), we hypothesize that the number of sites on the genome that dCas9 interacts with for longer than ~40ms is in the same order of magnitude as the highest dCas9 copy numbers (642). If many more sites on the genome would be potentially occupied by dCas9, there would be no relationship between dCas9 copy number and genomic bound dCas9 population.

We can further estimate the amount of partially complementary DNA sites by taking the size of the genome in *L. lactis* into account (2.5 million base pairs). Assuming that all bases occur with similar frequency, 2-3 sites in the complete genome will have 10-bp-sgRNA-DNA complementarity, which increases to ± 800 sites for 6+ bp-sgRNA-DNA complementarity and to ± 3,000 sites for 5+ bp-sgRNA-DNA complementarity. However, we do not know if the decrease in genomic-bound dCas9 continuous beyond 642 dCas9 copy numbers per cell, which could indicate that more partial target sites with high enough sgRNA-DNA complementarity for > 40ms dCas9 interaction are present on the genome. Moreover, this discussion does not take physical accessibility of the DNA into account.

### Supplementary Note 5: dCas9-PAmCherry2 DNA and AA sequences

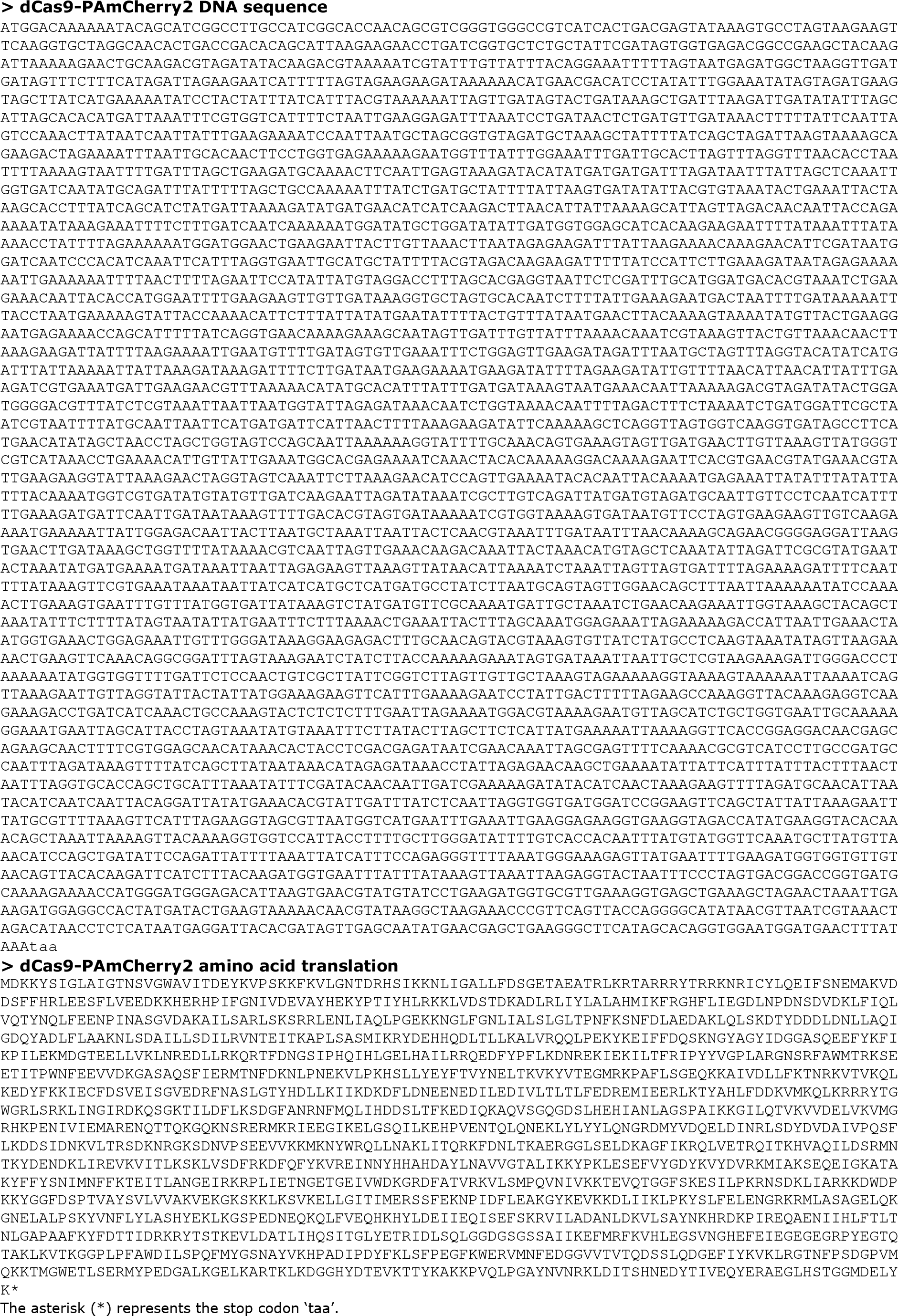

### Supplementary Note 6: Quantifying drift on a miCube

Linear drift calculations indicate that the system experiences a drift of 12.6 ± 11.6 nm/min in the lateral plane and 24.9 ± 14.5 nm/min in the axial plane. This is the average of three measurements performed on three different days (i.e. nine measurements in total). A typical drift measurement is shown in Supplementary Figure 9.

**Supplementary Figure 1:**
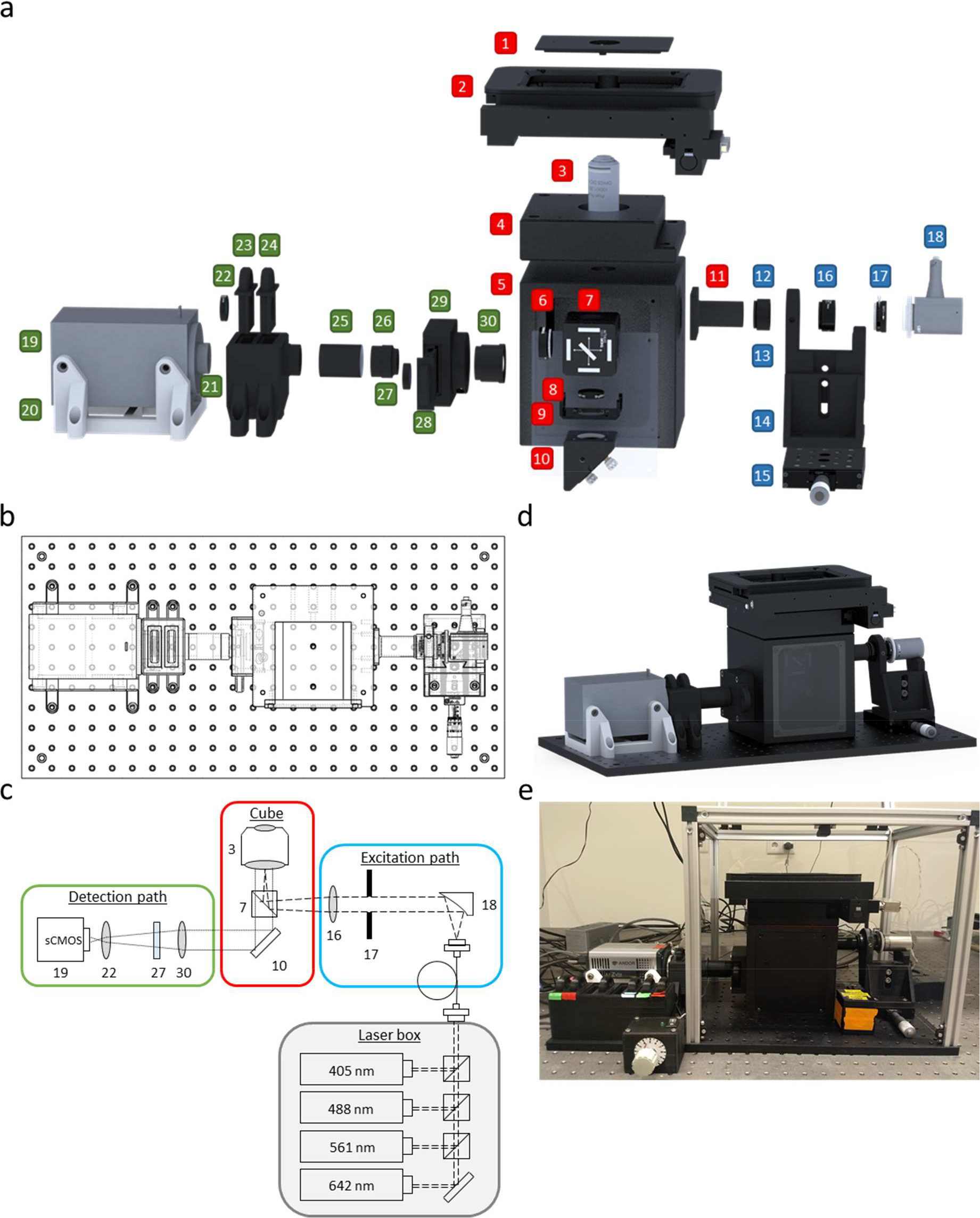
Overview of the miCube. (**a**) Exploded render of the miCube highlighting individual components. A full list of components indicated by the numbered items can be found in Supplementary Table 5. (**b**) Top-down schematic view of the miCube on the breadboard, allowing clear view of mounting positions. Distance between mounting holes on the breadboard is 25 mm. (**c**) Schematic overview of the miCube instrument. Numbered items correspond to the items in (a) and Supplementary Table 5. The excitation path is visualized with dashed lines, the emission path is visualized with dotted lines. (**d**) Virtual rendition of the fully assembled miCube instrument placed on a Thorlabs breadboard. (**e**) Photograph of the fully assembled miCube as used for measurements in this manuscript.

**Supplementary Figure 2:**
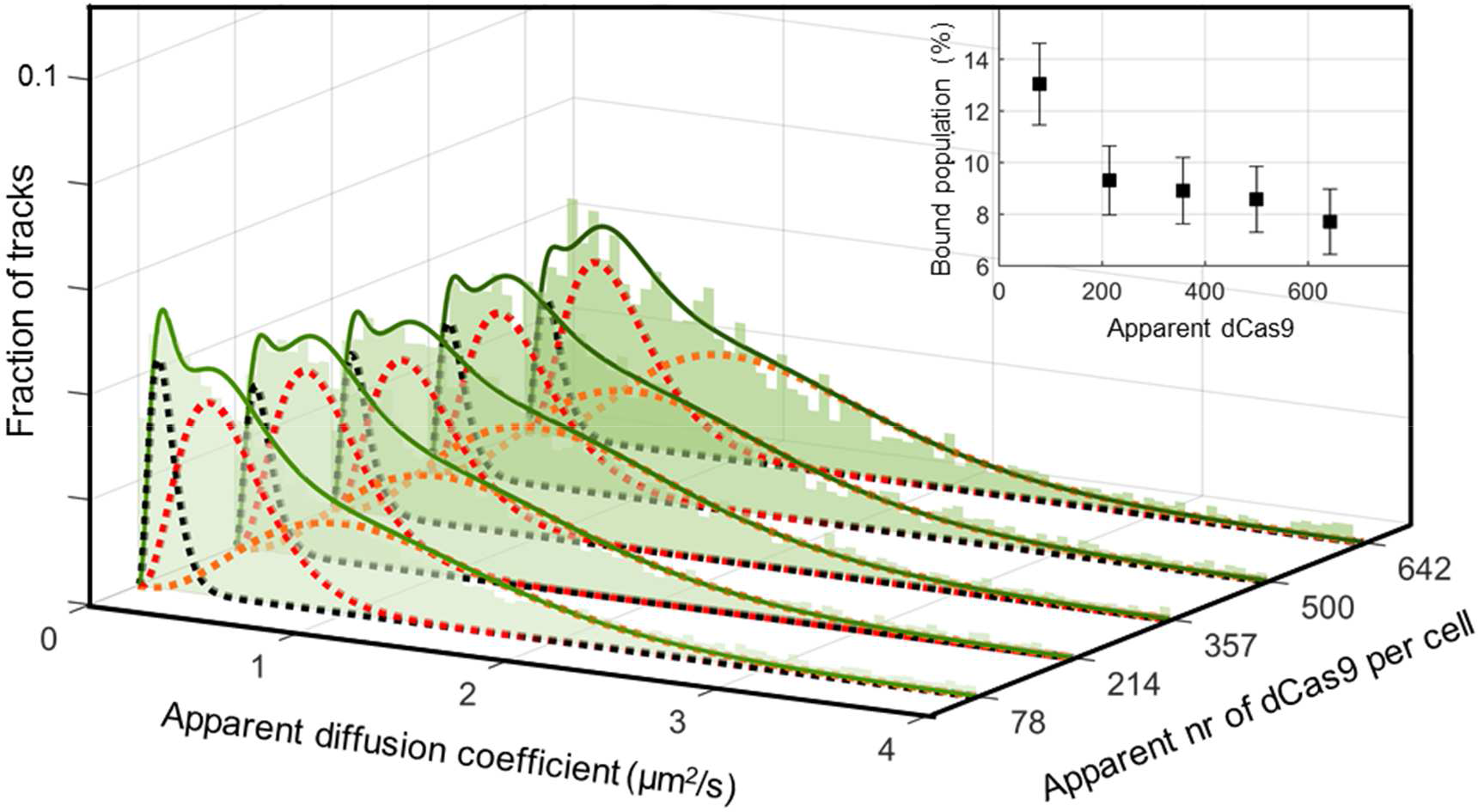
dCas9 diffusion in cells containing pNonTarget. The apparent diffusion coefficient histograms (bin width = 0.05 µm^2^/s) of pNonTarget-containing cells were fitted with populations corresponding to 0.13 (black), 0.46 (red), and 1.31 (orange) µm^2^/s, after being corrected for *L. lactis* cells lacking the dCas9-sgRNA plasmid. The inset shows the bound population as function of apparent number of dCas9 per cell corrected for *L. lactis* cells lacking the dCas9-sgRNA plasmid. The error bar indicates the 95% confidence interval of the fit of the bound population with the data (least-squares). Uncorrected populations are used as partial fit of the apparent diffusion histograms arising from pTarget-containing cells (Fig. 1d).

**Supplementary Figure 3:**
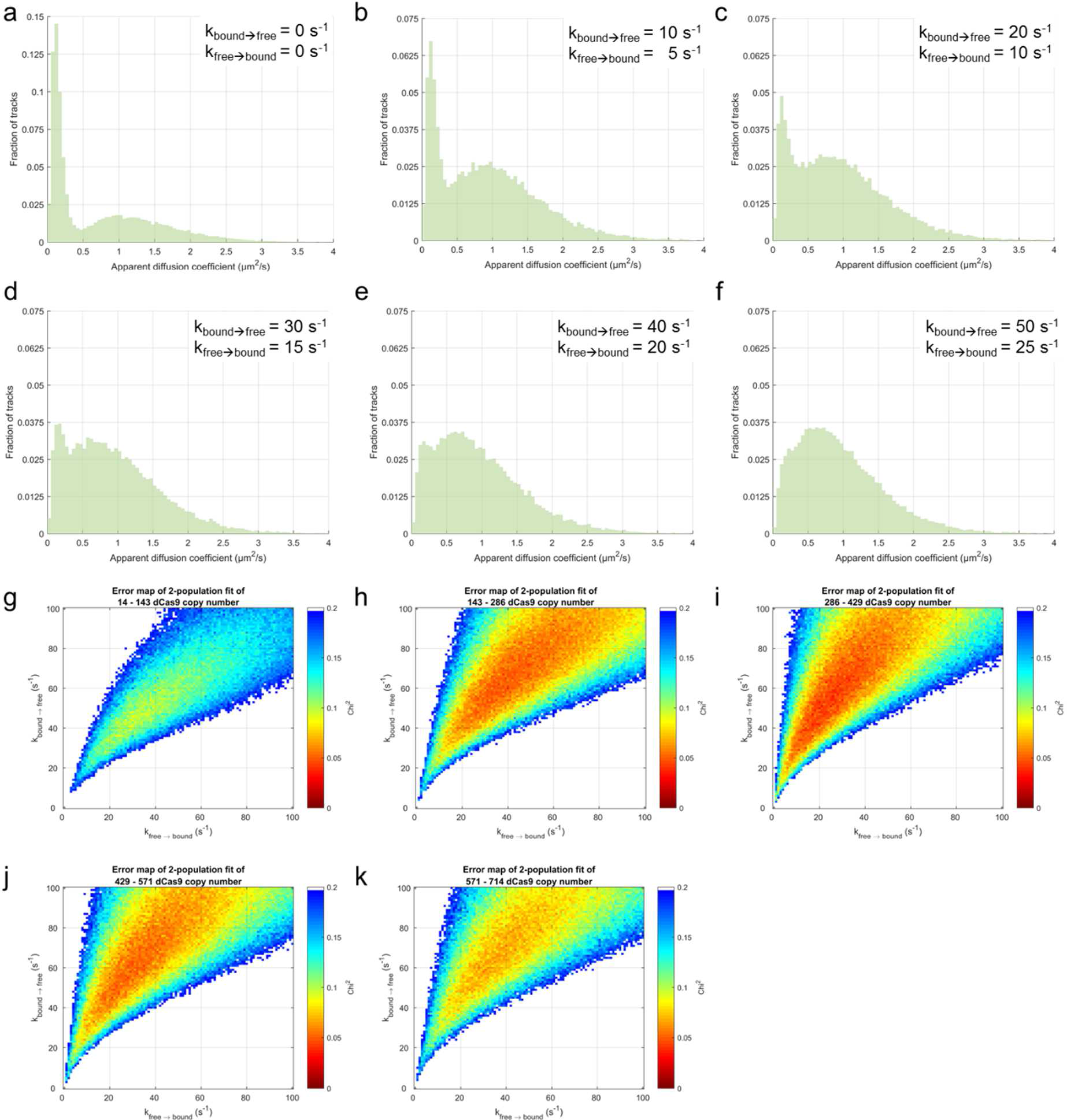
Temporal averaging due to fast switching of diffusion coefficients. A simulation was run with two populations (*D*_bound_ and *D*_free_), while each simulated molecule was free to switch between the two populations according to the rate constants *k*_bound→free_ and *k*_free→bound_. (**a-f**): Simulated diffusion coefficient histograms with increasing rate constants, but constant ratio between the rate constants. (**g-k**): Error maps showing the *Χ*^2^ error between the pNonTarget diffusion coefficient histograms with different dCas9 copy number interval (indicated in title) and simulations with different rate constants. A full description of the simulations can be found in Supplementary Note 3.

**Supplementary Figure 4:**
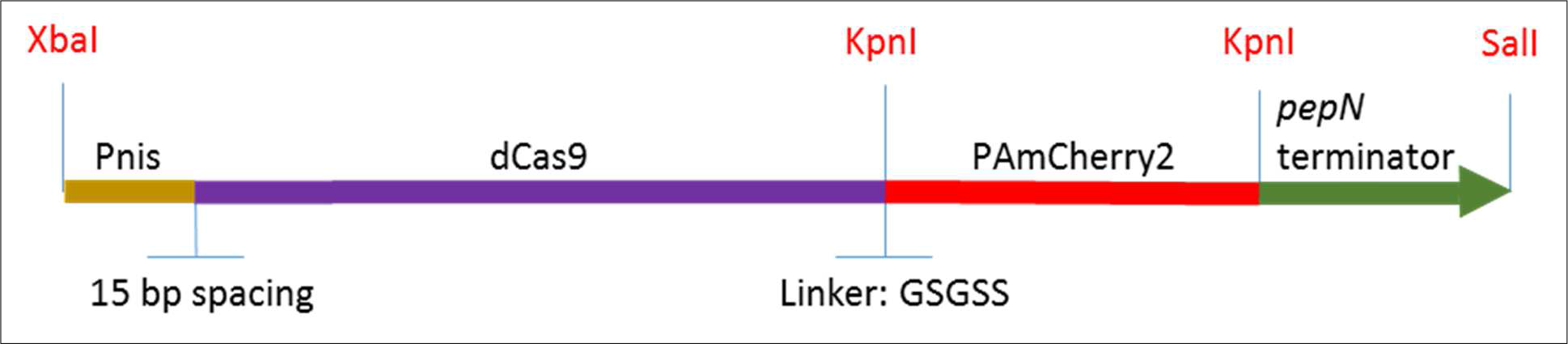
Genetic design of Pnis-dCas9-PAmCherry2. The sequence encoding dCas9^33^ (*S. pyogenes*; AddGene.org plasmid #44249) is fused to the sequence encoding PAmCherry2^28^ (AddGene.org plasmid #31932) with a flexible linker (amino acid sequence GSGSS), downstream of the *nisA*-promoter (Pnis) with an ribosomal binding site (15 bp spacing) and ending with a transcriptional terminator sequence derived from a lactococcal *pepN* gene. PAmCherry2 is flanked by two KpnI sites which should allow for interchanging fluorophores. The whole sequence is flanked by XbaI and SalI restriction sites to allow convenient cloning into a (expression) vector of choice.

**Supplementary Figure 5:**
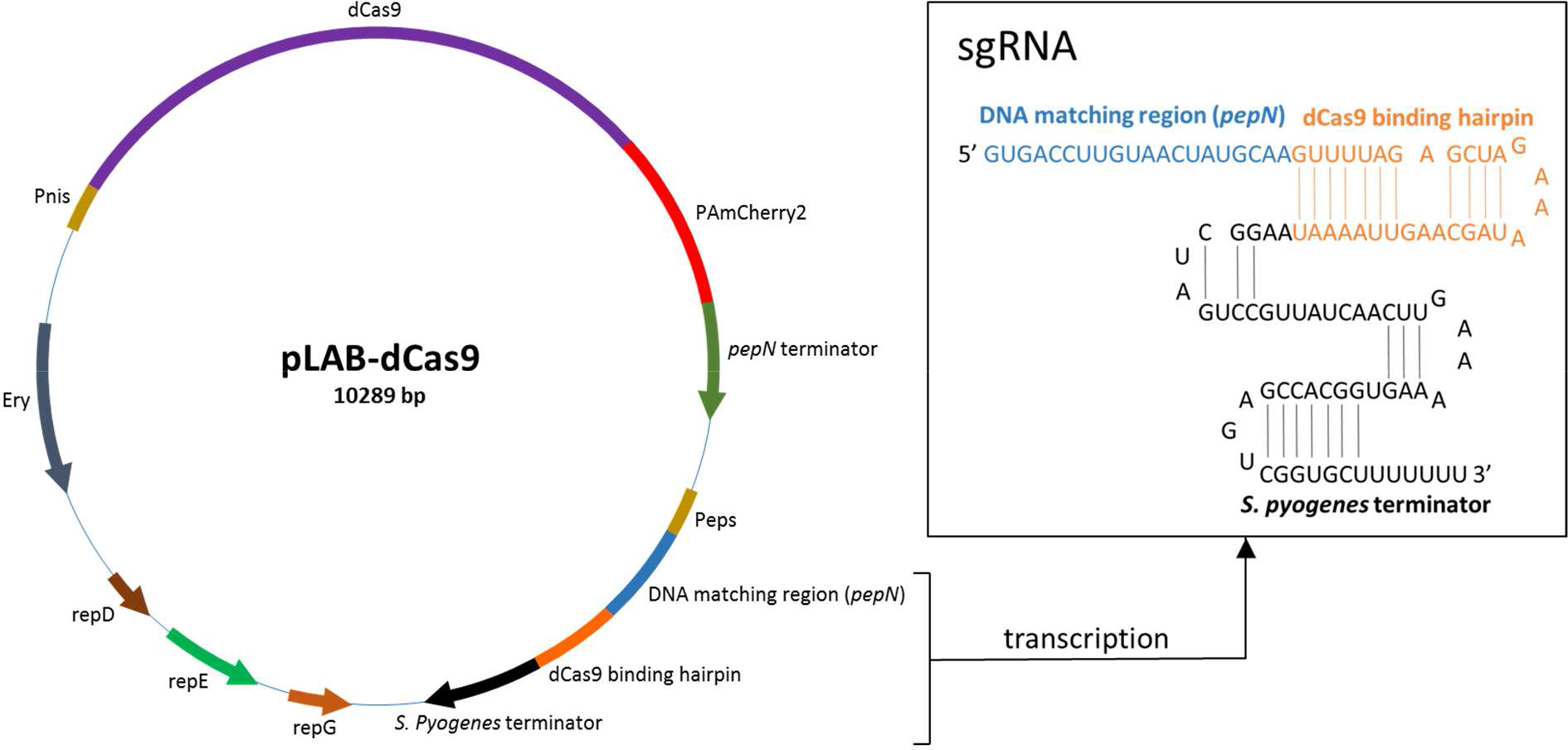
Outline of the pLAB-dCas9 vector. The pLAB-dCas9 expression vector consists of nisin-inducible (Pnis) PAmCherry2-labelled dCas9, an erythromycin resistance marker (Ery) and replication genes (*repD*, *repE* and *repG*)^34^. The *pepN* DNA matching region together with the dCas9 binding hairpin and the *S. pyogenes* terminator form the sgRNA, which is expressed under a constitutive promoter (Peps). Once the sgRNA molecule is transcribed, it folds to form the secondary structure that allows complex formation with dCas9.

**Supplementary Figure 6:**
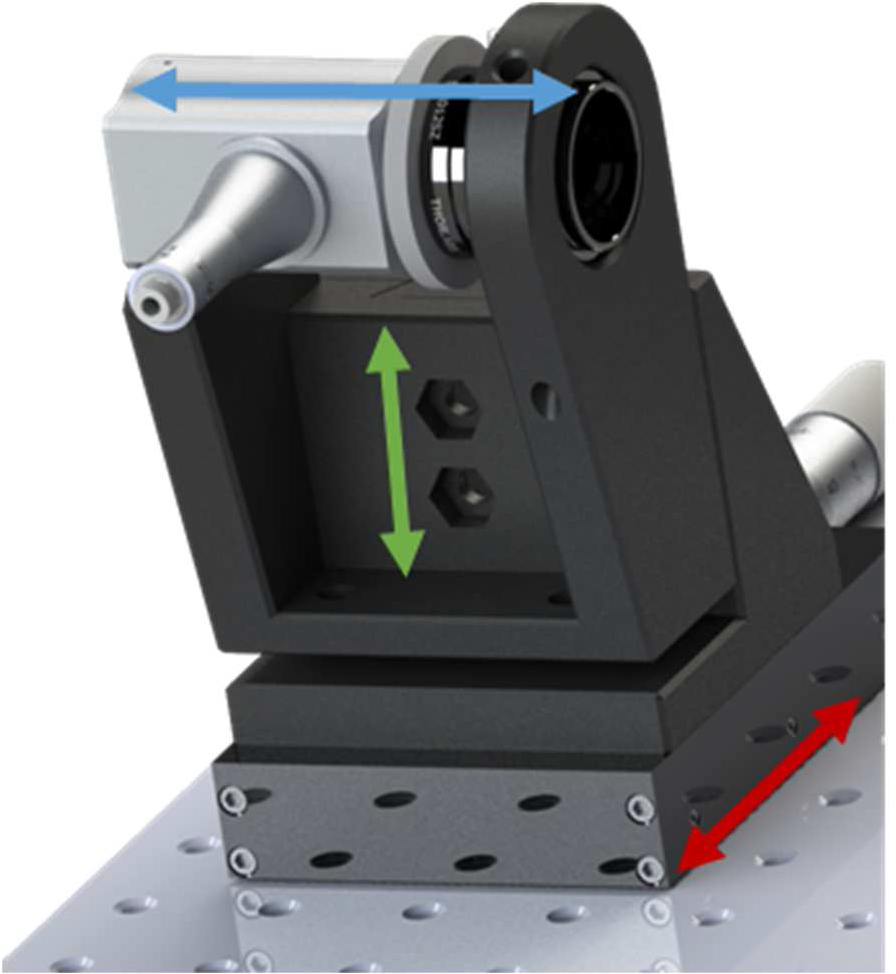
Detailed view of the excitation path. This sub-part is comprised of numbers 12-18. Arrows indicate isolated movement in all three dimensions: distance from objective (blue), height of excitation unit (green), and horizontal position with respect to the objective (red).

**Supplementary Figure 7:**
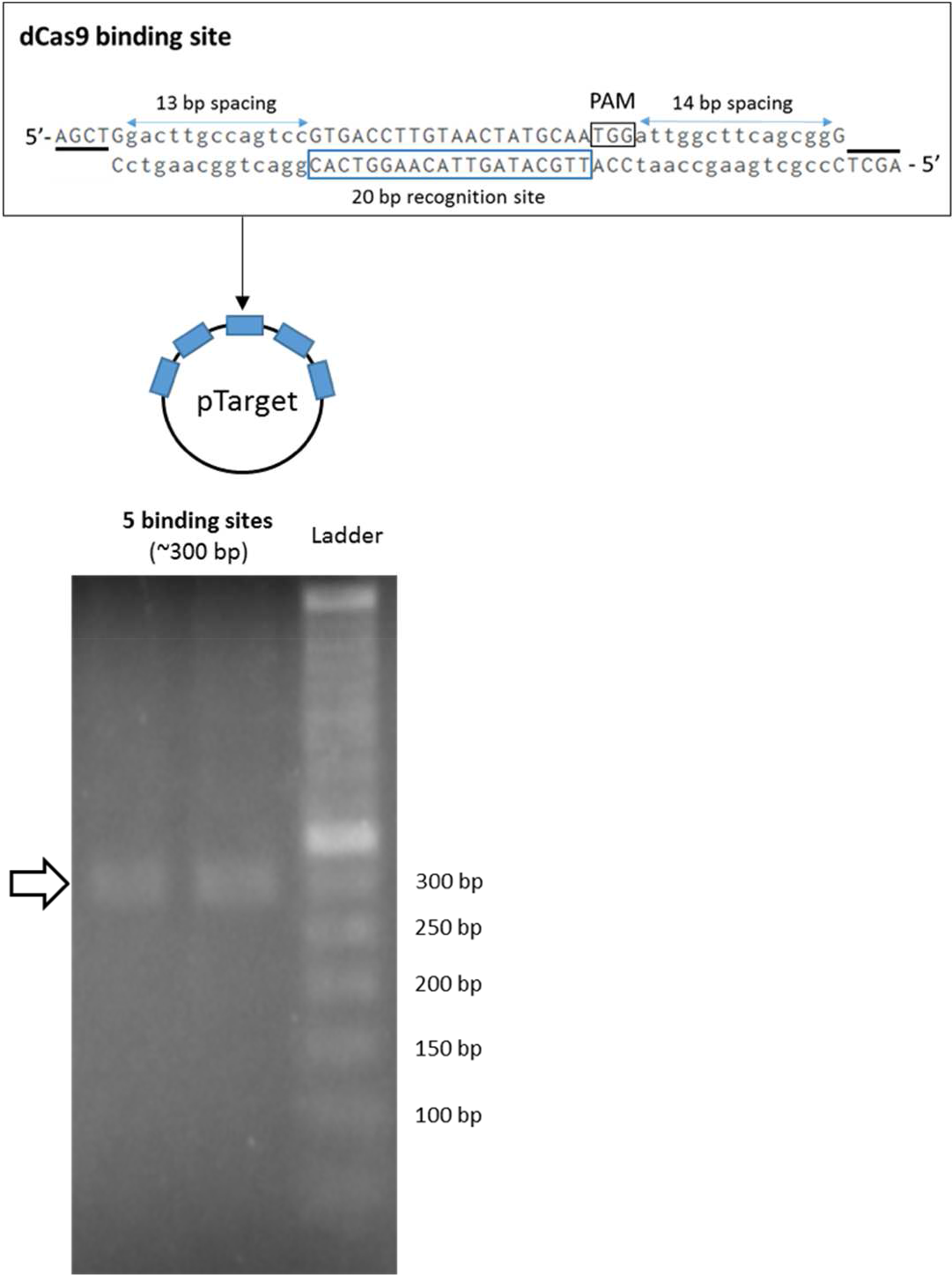
Construction and verification of pTarget. dCas9 binding sites consisting of a 20 base pairs *pepN* recognition site, a 5’-NGG-3’ PAM sequence, and spacing and overhang sequence motifs that are complementary to each other (indicated with black stripes) were annealed and ligated. This formed an array of five dCas9 binding sites in pNZ123, resulting in pTarget. Digestion and subsequent gel electrophoresis of plasmids isolated from two colonies revealed the expected length of the binding array in pTarget. Since one binding site is 60 base pairs in length, an array of five binding sites is ∼300 base pairs.

**Supplementary Figure 8:**
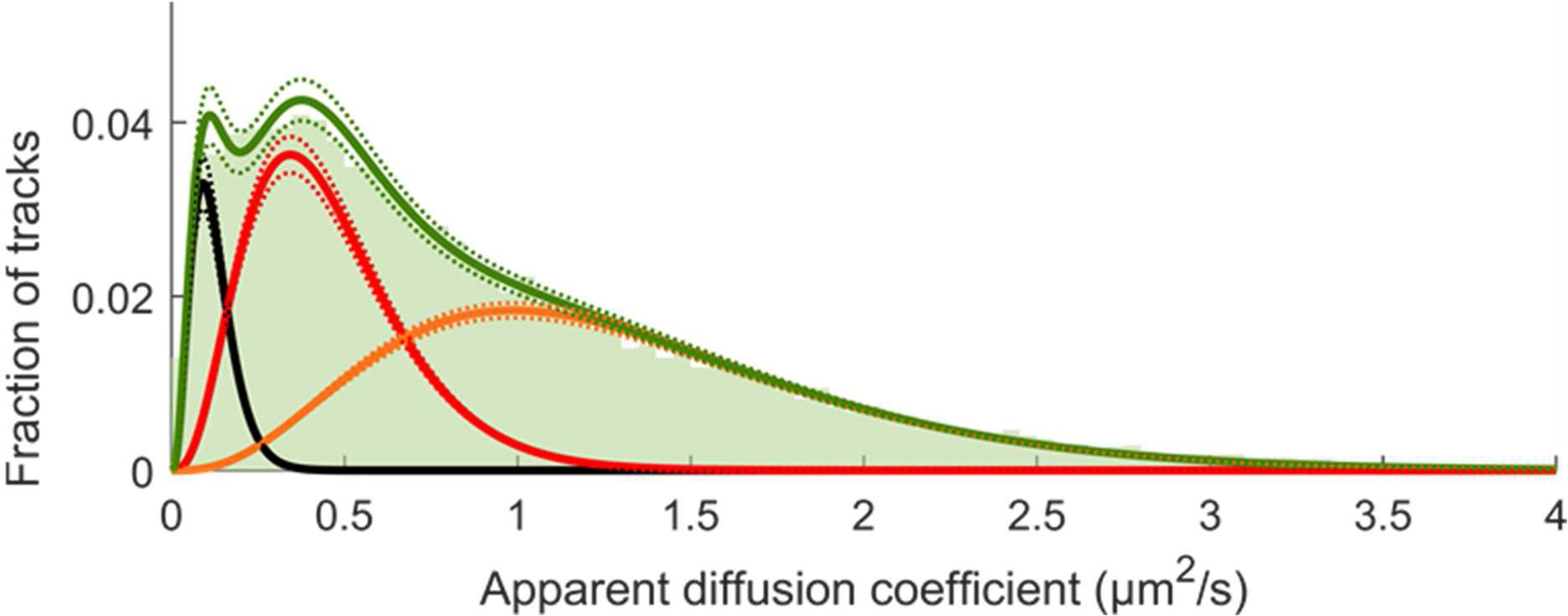
Histogram of simulated data (Supplementary Note 3). The simulated data contained 33.000 tracks of particles with real diffusion coefficients of 0 µm^2^/s, 0.43 µm^2^/s and 1.95 µm^2^/s in a 8:35:57 ratio, with an added 36.5 nm localization uncertainty, in a modelled cell consisting of 2 half-spheres with radius 0.5 µm connected by a straight cylinder of length 0.5 µm and radius 0.5 µm. See Supplementary Note 3 for more details concerning the simulation. The resulting parameters are apparent diffusion coefficients of 0.12-0.13 µm^2^/s, 0.44-0.48 µm^2^/s, and 1.26-1.36 µm^2^/s (black, red, and orange, respectively) in a 9-10:44-48:51-56 ratio. This is similar to the experimental data fitted in Figure 1c.

**Supplementary Figure 9:**
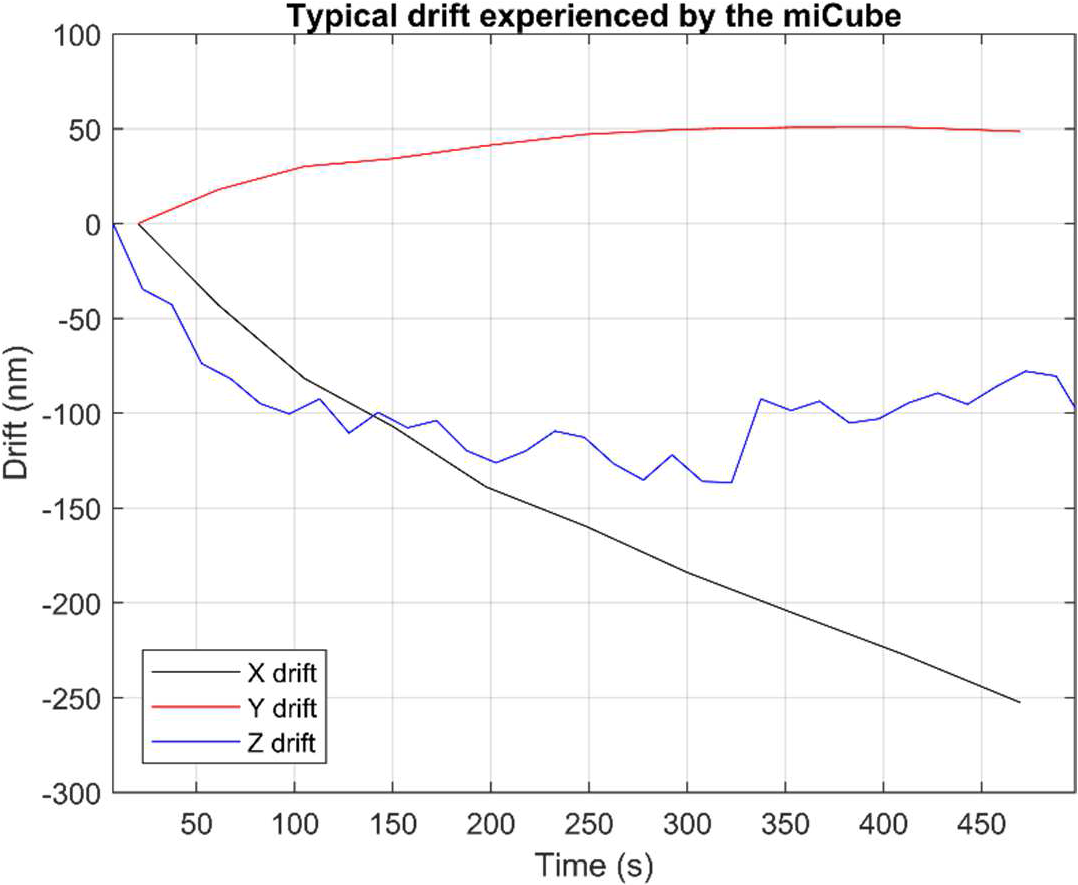
Typical drift experienced by the miCube. Typical drift in X (black), Y (red), and Z (blue) as experienced by the miCube used throughout this study. Repetition of this experiment led to the values specified in Supplementary Note 6.

**Supplementary Table 1:** List of vectors.

| Vector | Relevant properties | Size (kb) | Selection marker | Reference |
| --- | --- | --- | --- | --- |
| pNZ123 | High-copy number (~60) vector replicating in *L. lactis*. | 2.8 | Chloramphenicol | <sup>35</sup> |
| pLABTarget | Encoding functional Cas9 expression system | 10 | Erythromycin | <sup>4</sup> |
| pNonTarget | pNZ123 without binding sites | 2.8 | Chloramphenicol | This study |
| pTarget | pNZ123 containing five binding sites cognate to *pepN* sgRNA | 3.1 | Chloramphenicol | This study |
| pLAB-dCas9-nosgRNA | Cas9 module of pLABTarget replaced with Pnis – dCas9 – PAmCherry2 | 10.2 | Erythromycin | This study |
| pLAB-dCas9 | pLAB-dCas9-nosgRNA with added *pepN* sgRNA under constitutive promotor | 10.3 | Erythromycin | This study |

**Supplementary Table 2:**
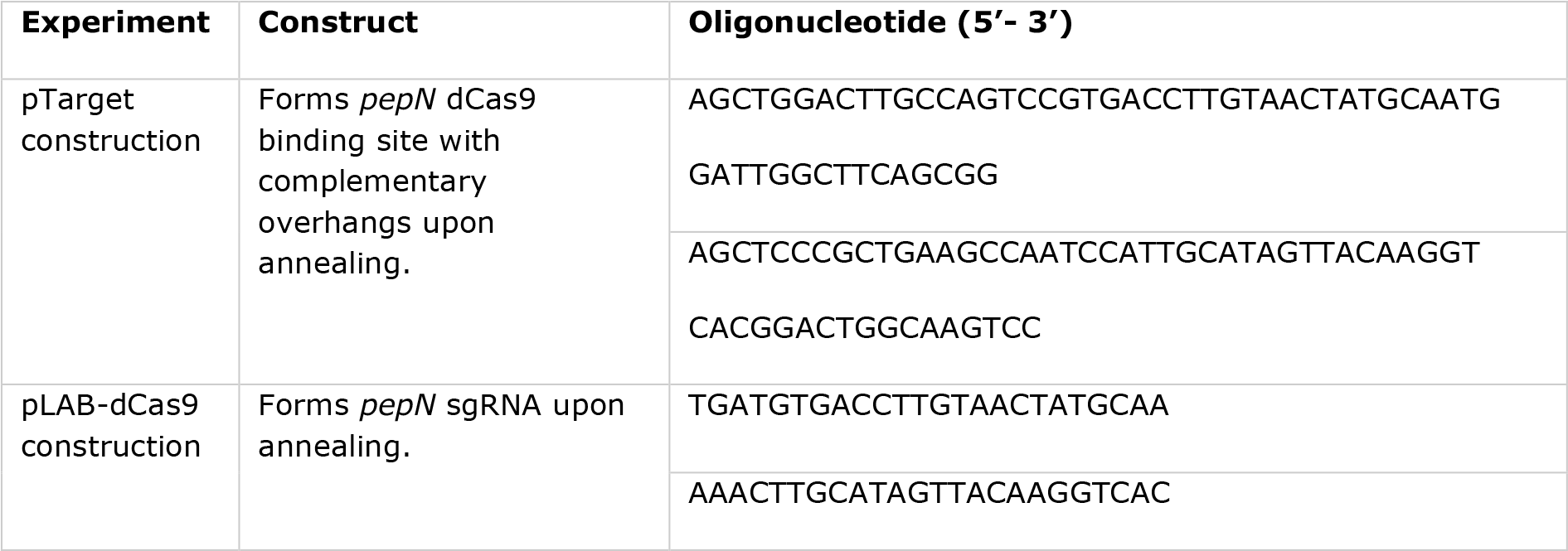
List of oligonucleotides.

**Supplementary Table 3:**
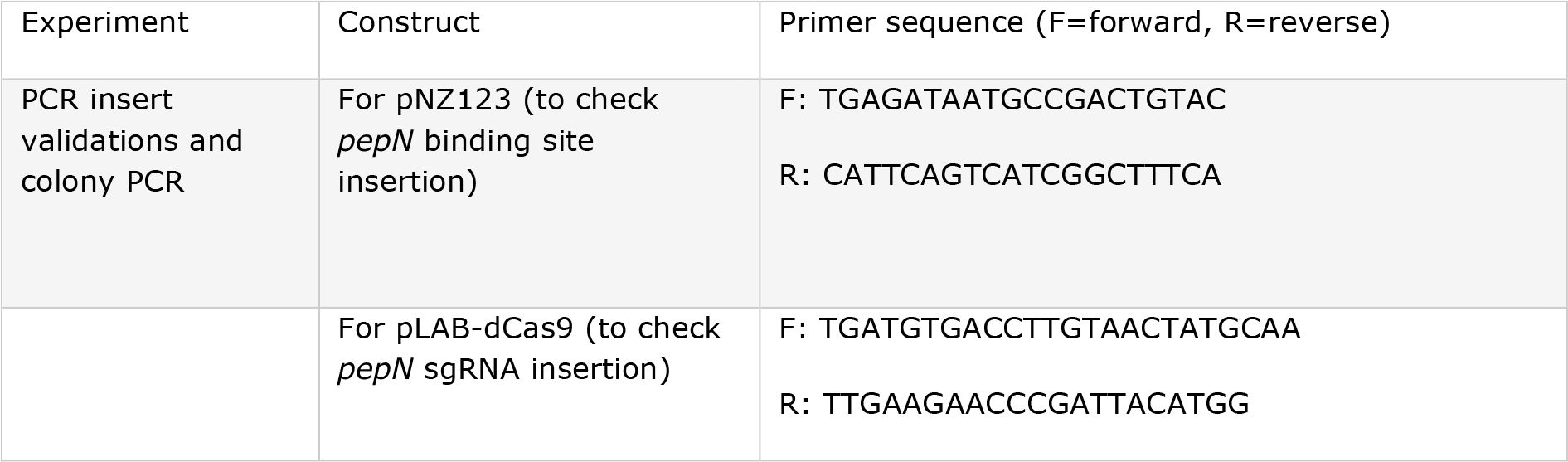
List of primers.

**Supplementary Table 4:**
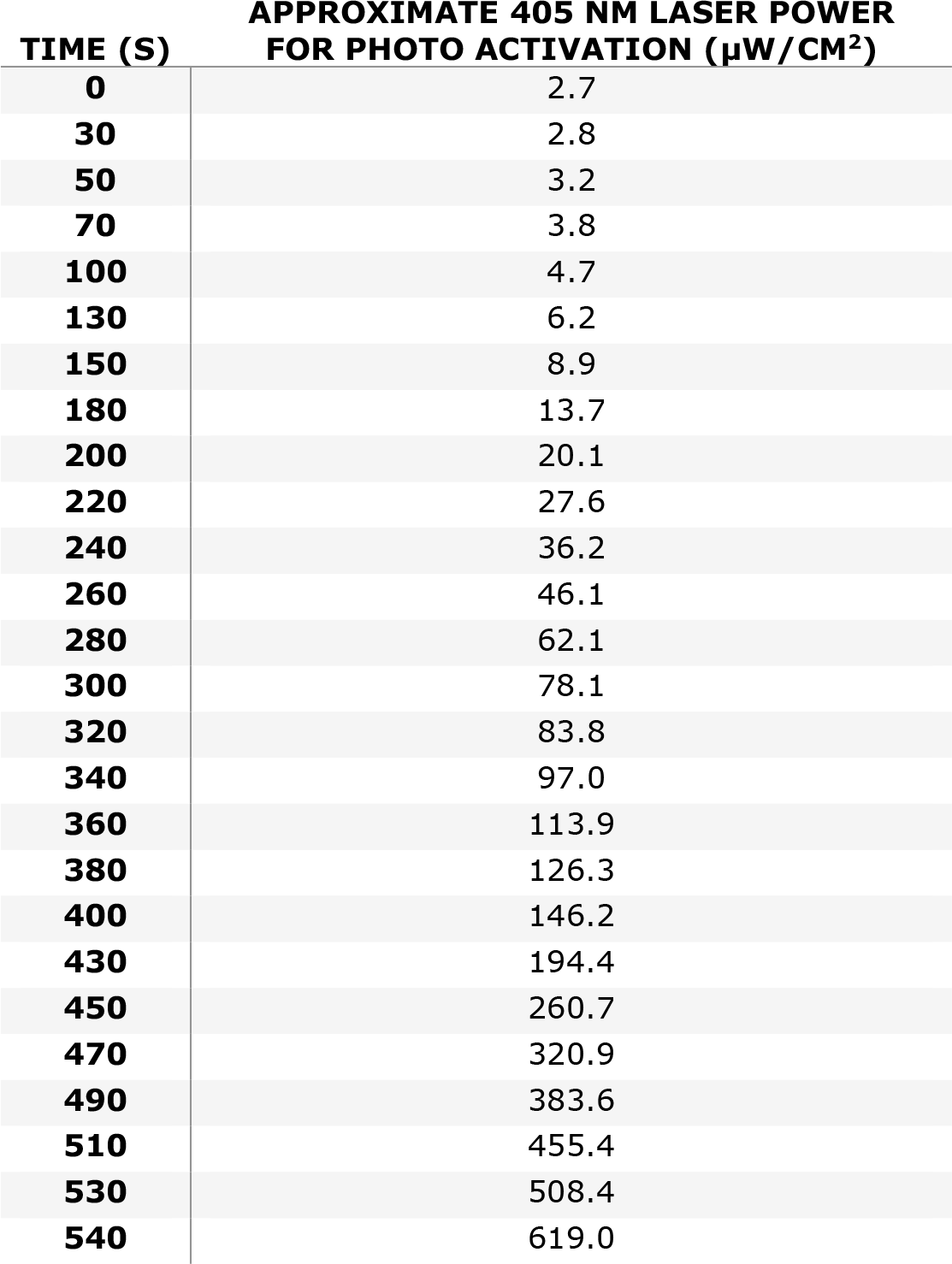
Adjustment of the 405 nm laser power during sptPALM experiments.

| **TIME (S)** | **APPROXIMATE 405 NM LASER POWER**<br>**FOR PHOTO ACTIVATION ($\mu\text{W}/\text{CM}^2$)** |
| --- | --- |
| **0** | 2.7 |
| **30** | 2.8 |
| **50** | 3.2 |
| **70** | 3.8 |
| **100** | 4.7 |
| **130** | 6.2 |
| **150** | 8.9 |
| **180** | 13.7 |
| **200** | 20.1 |
| **220** | 27.6 |
| **240** | 36.2 |
| **260** | 46.1 |
| **280** | 62.1 |
| **300** | 78.1 |
| **320** | 83.8 |
| **340** | 97.0 |
| **360** | 113.9 |
| **380** | 126.3 |
| **400** | 146.2 |
| **430** | 194.4 |
| **450** | 260.7 |
| **470** | 320.9 |
| **490** | 383.6 |
| **510** | 455.4 |
| **530** | 508.4 |
| **540** | 619.0 |

**Supplementary Table 5:**
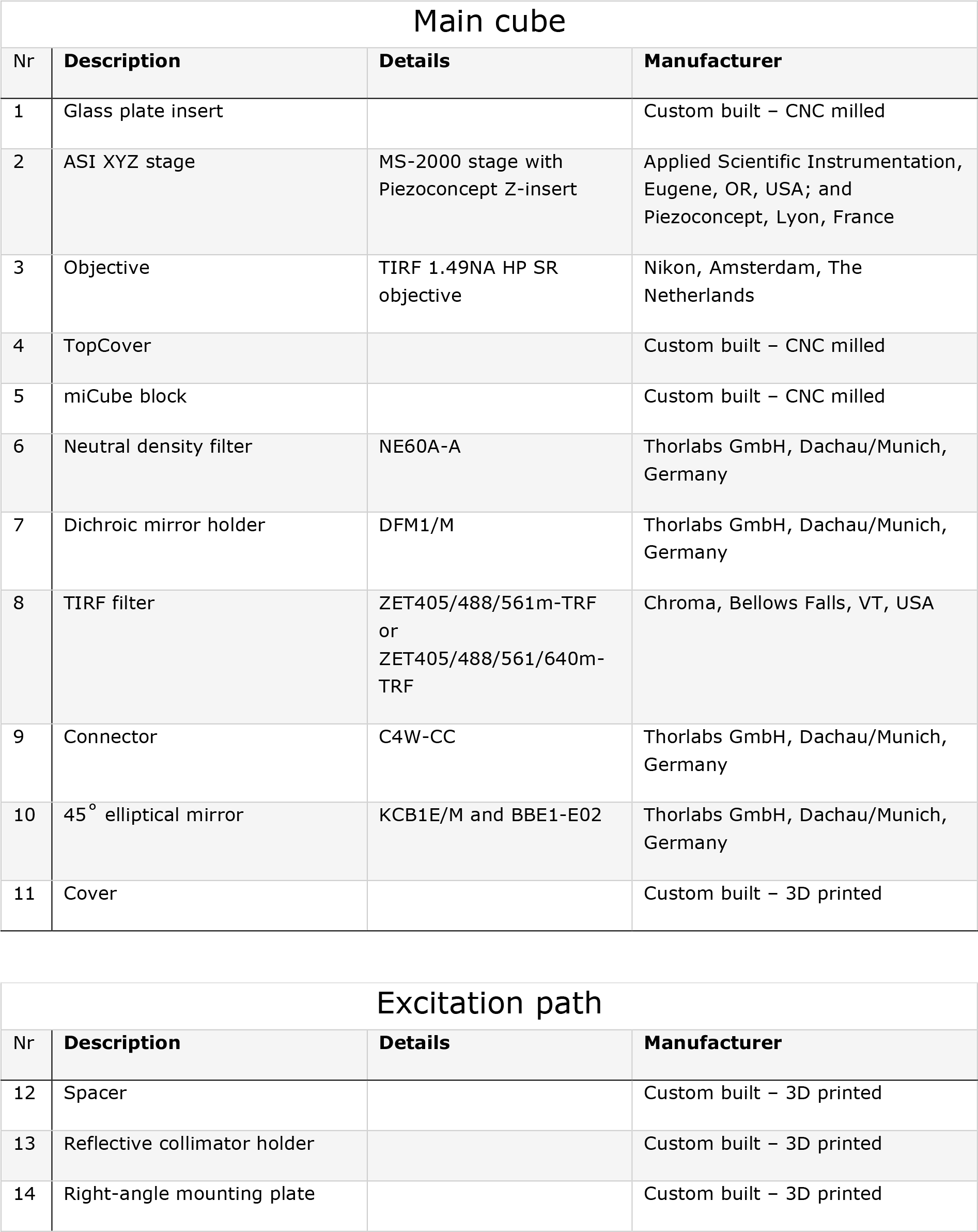
Descriptive list of components miCube. Numbers are in accordance with Supplementary Figure 1. Entities marked with ‘custom-built’ have their complete technical drawings present in the Appendix: Technical drawings of miCube components. A more exhaustive list can be found on https://HohlbeinLab.github.io/miCube/component_table.html.

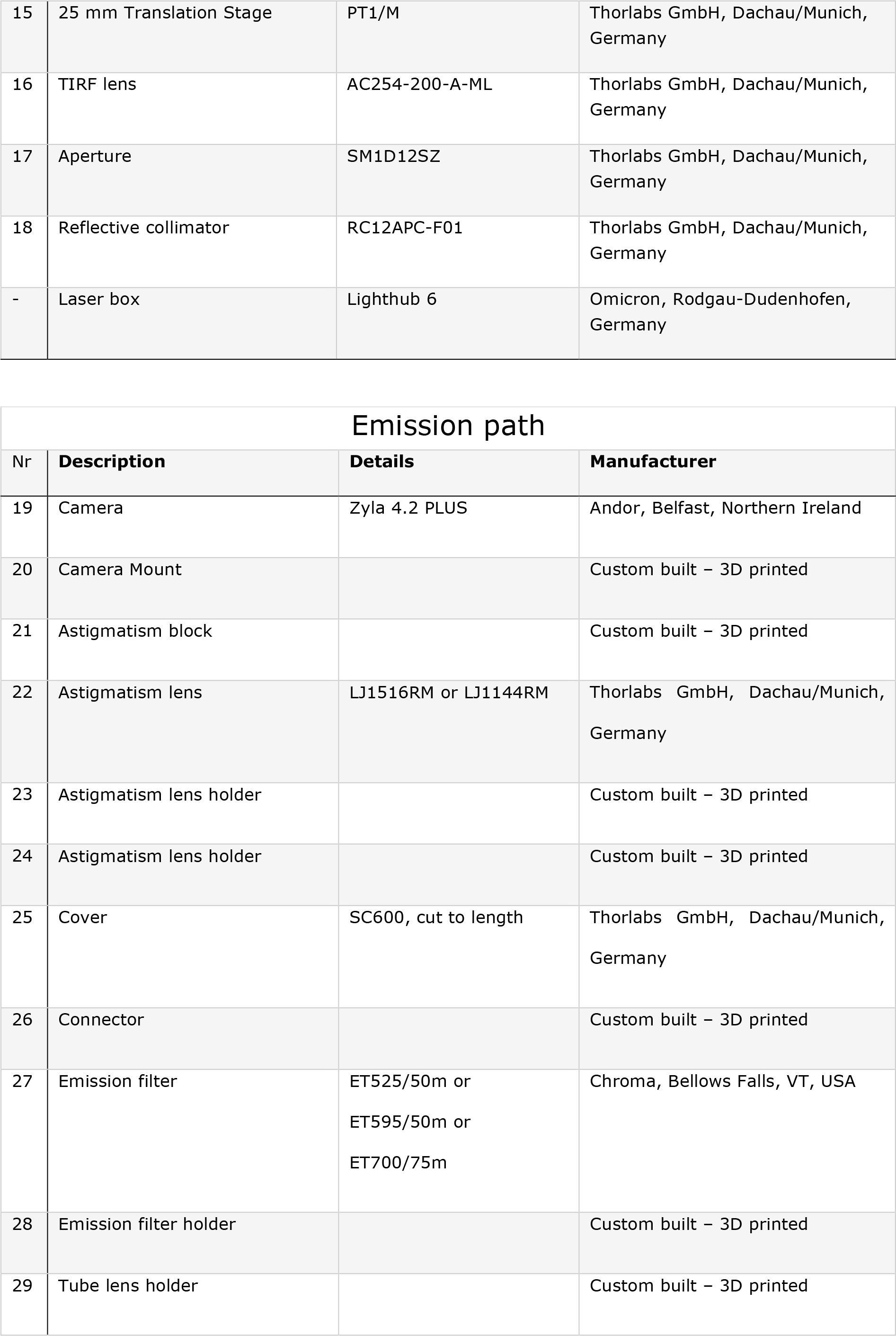

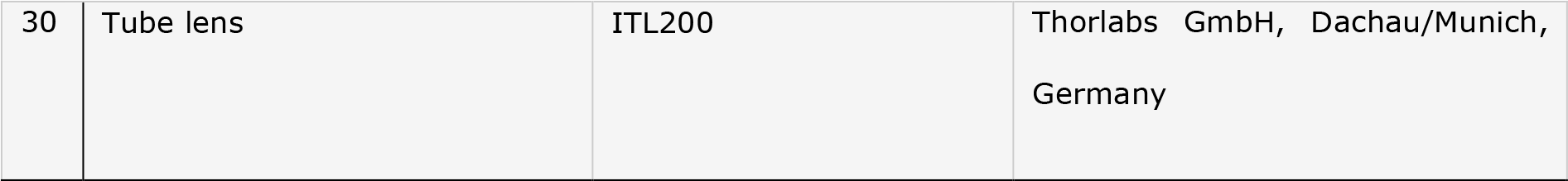

## Appendix: Technical drawings of miCube components

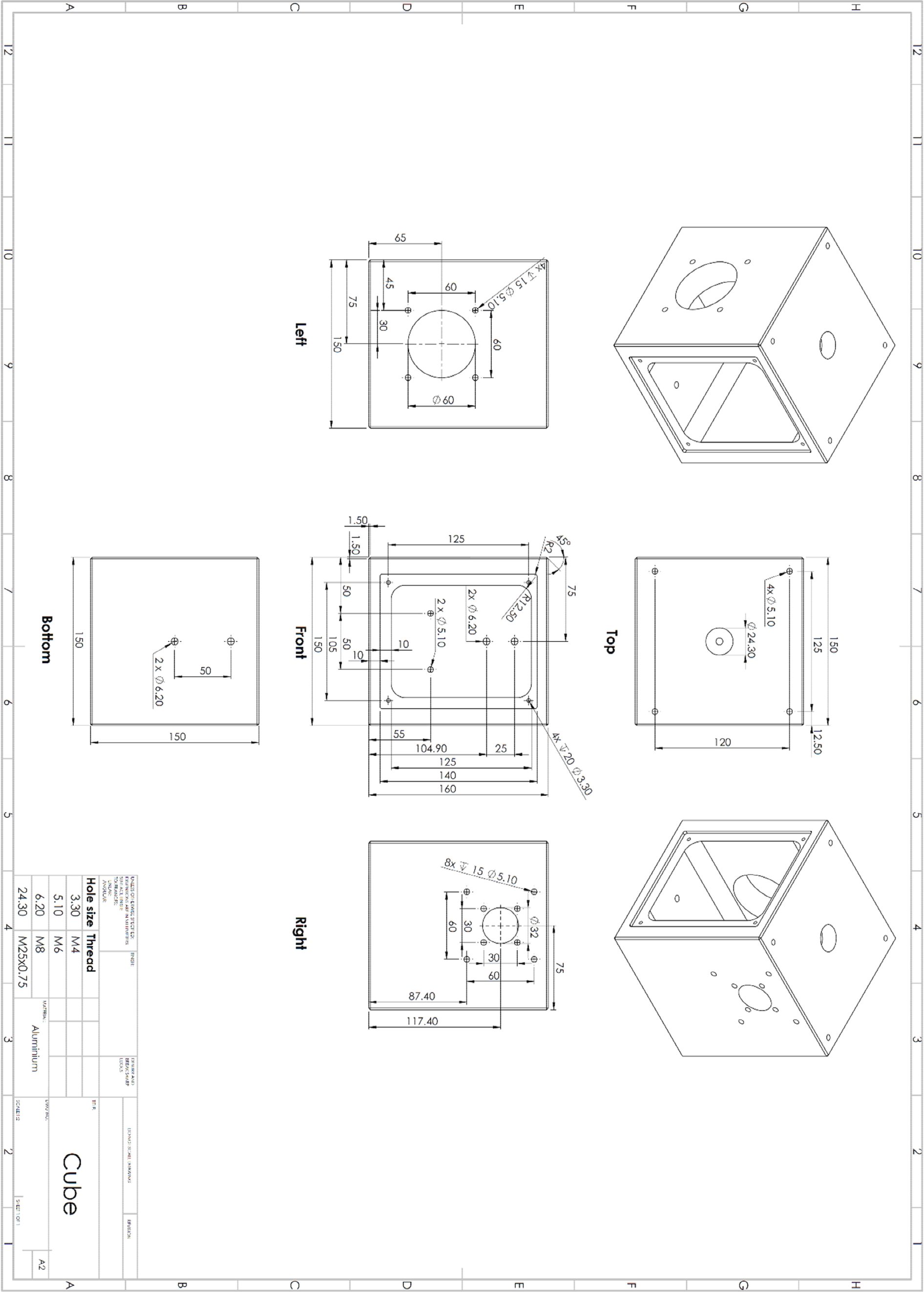

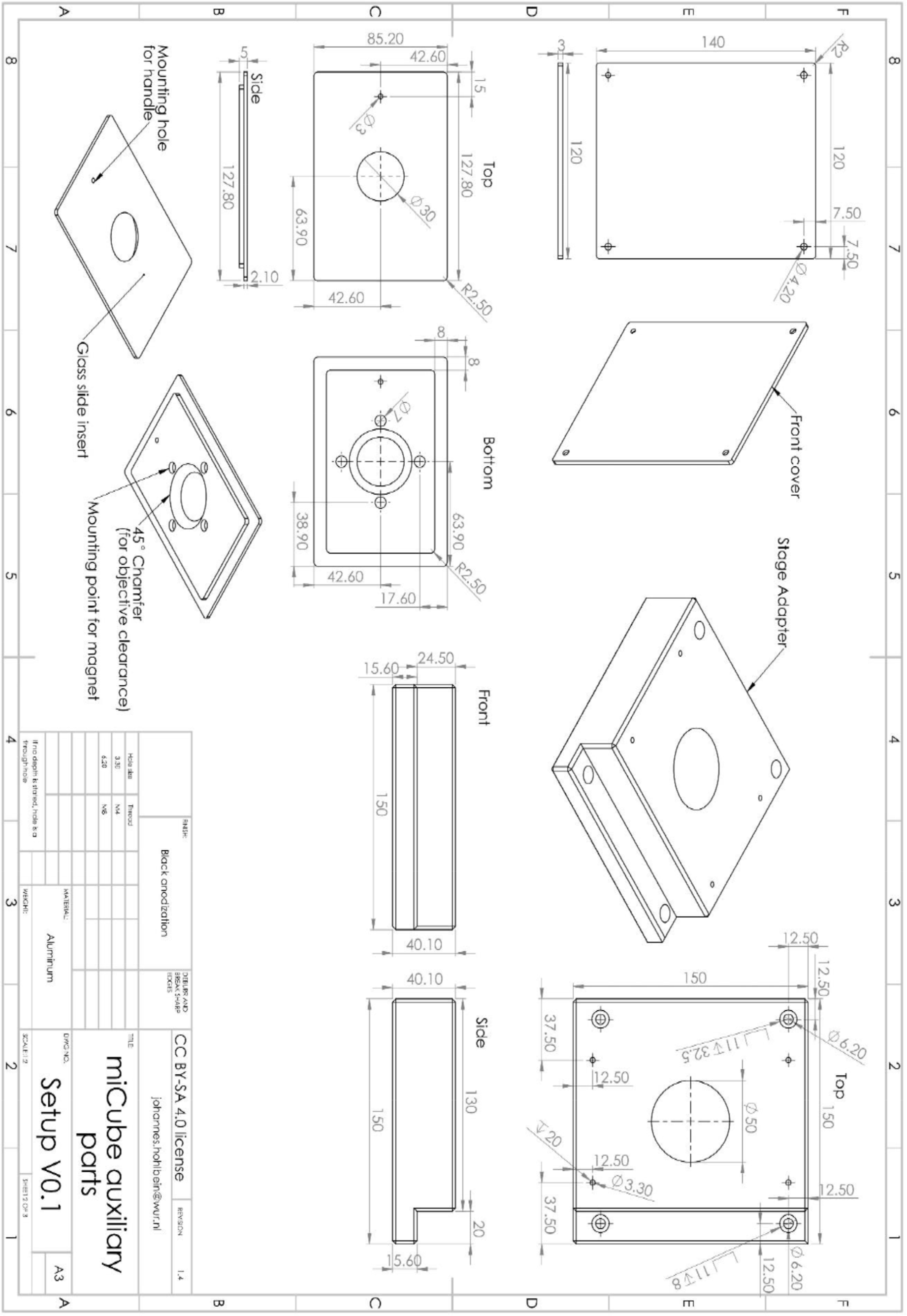

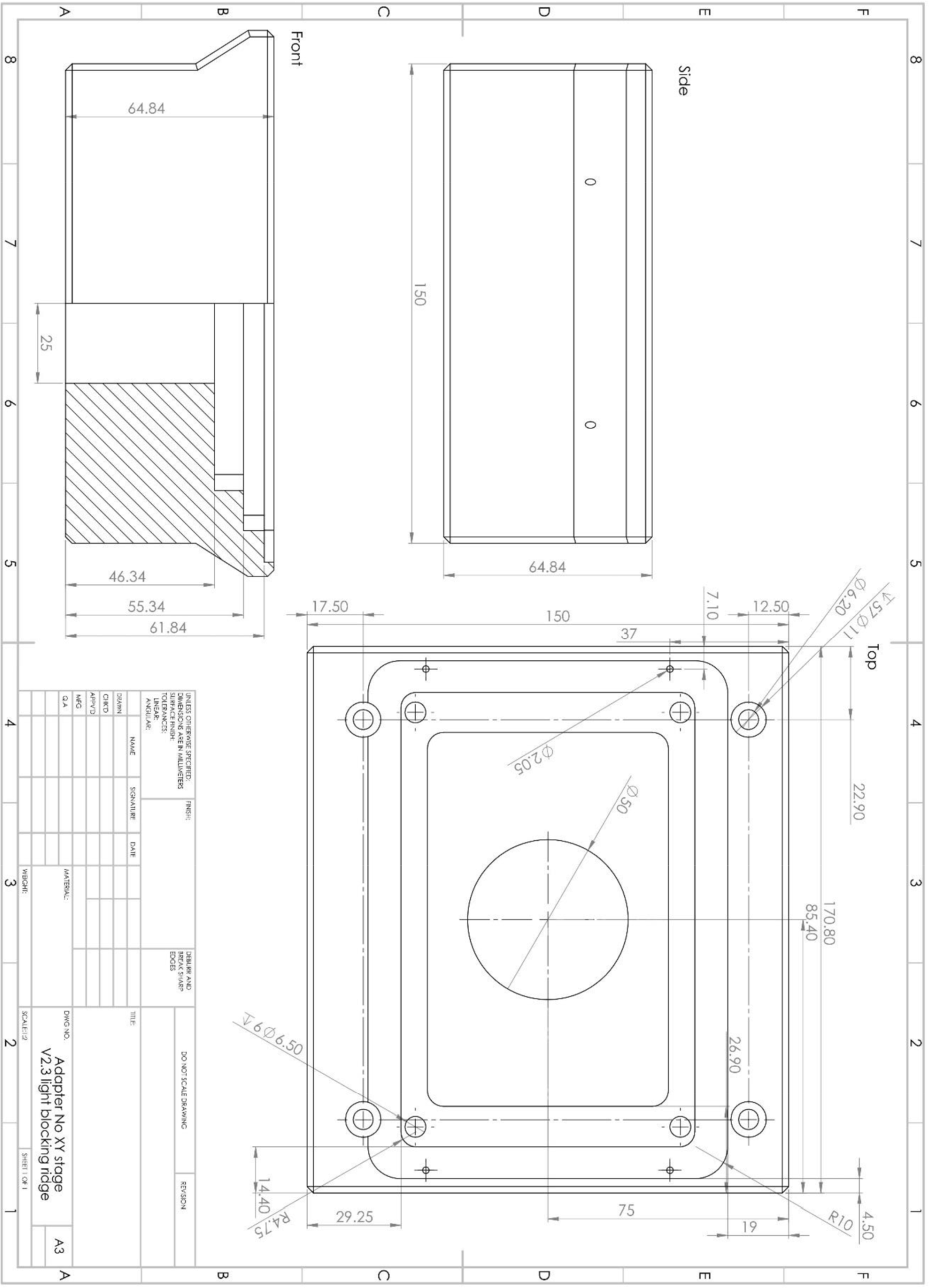

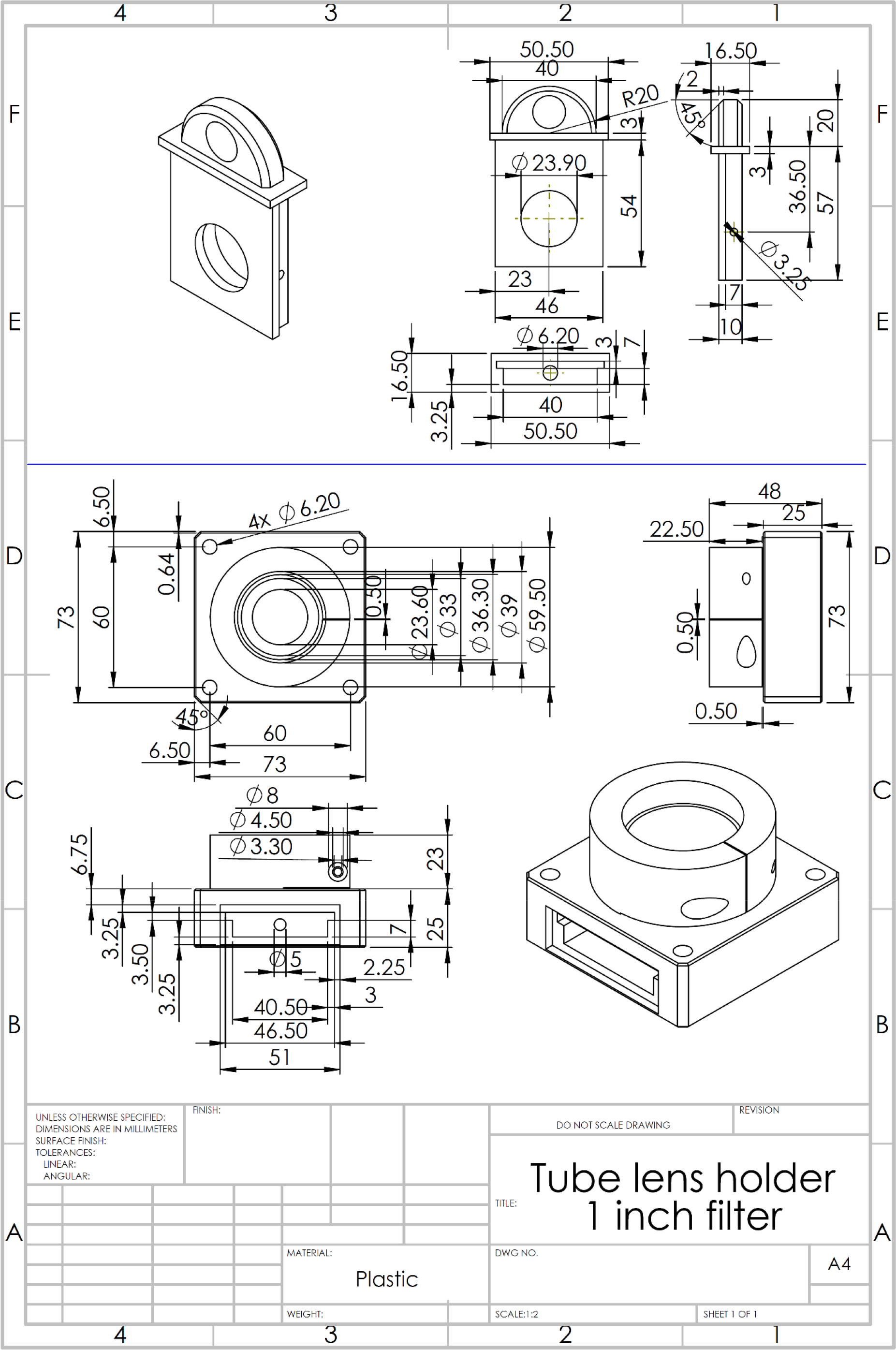

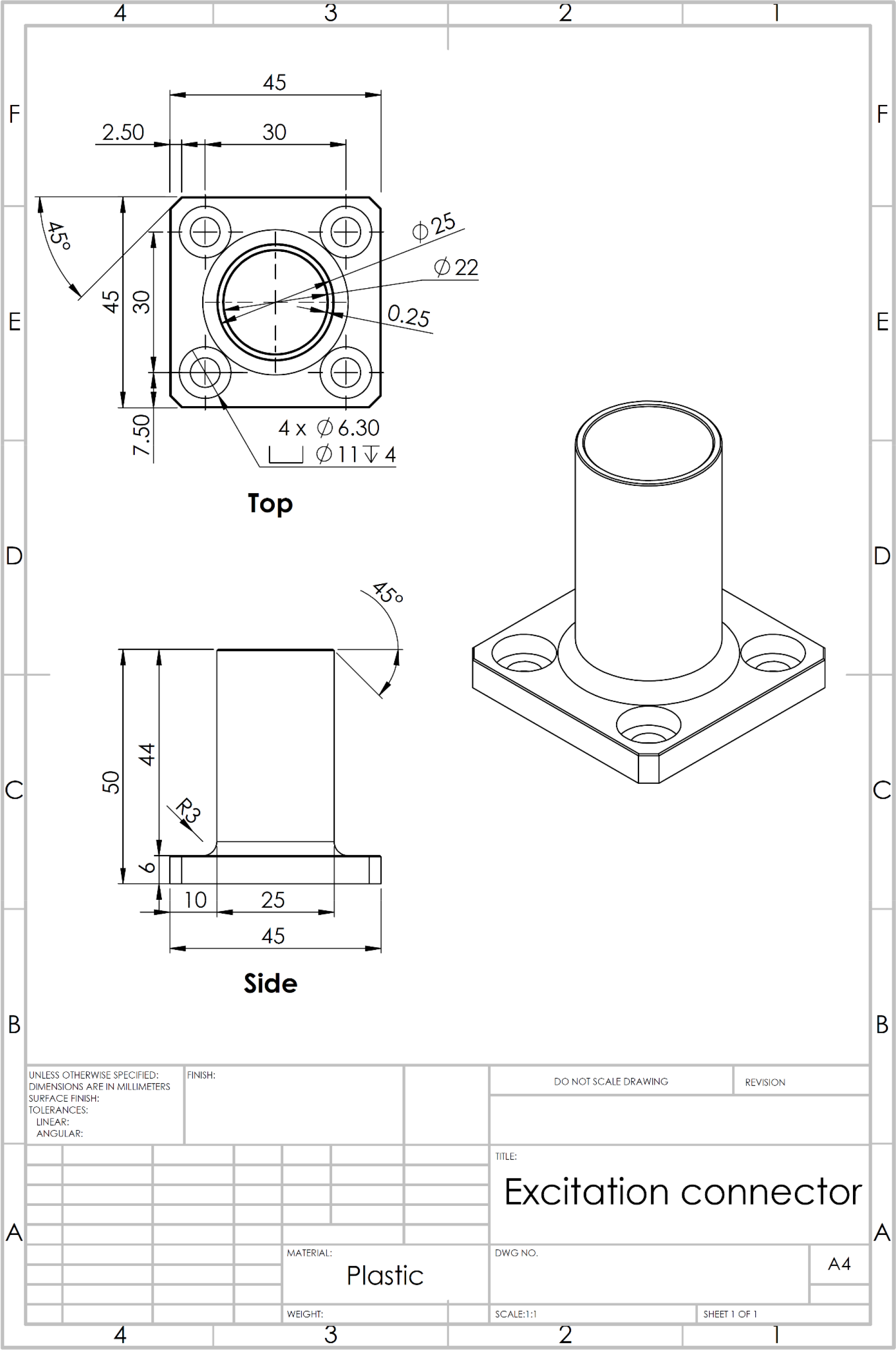

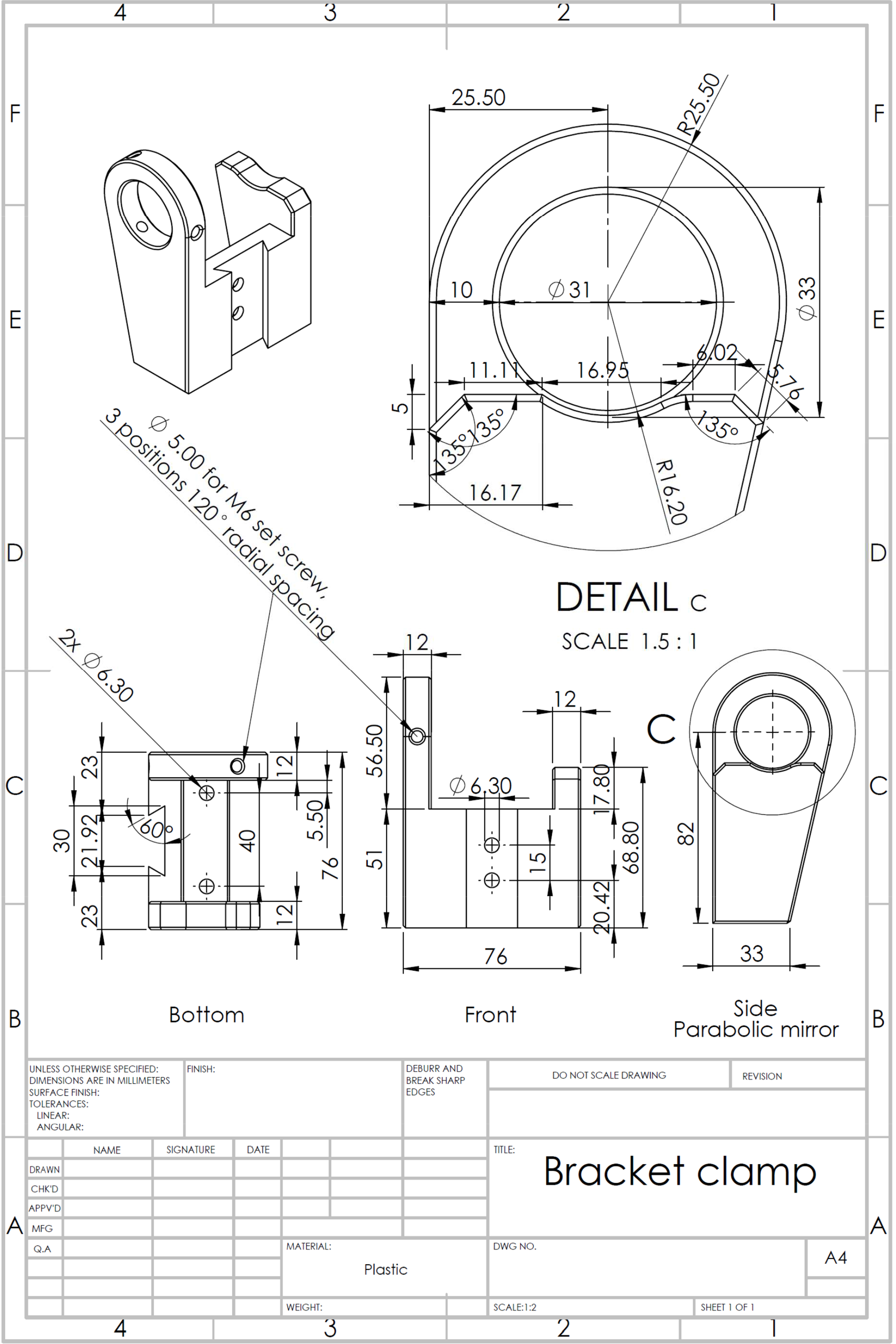

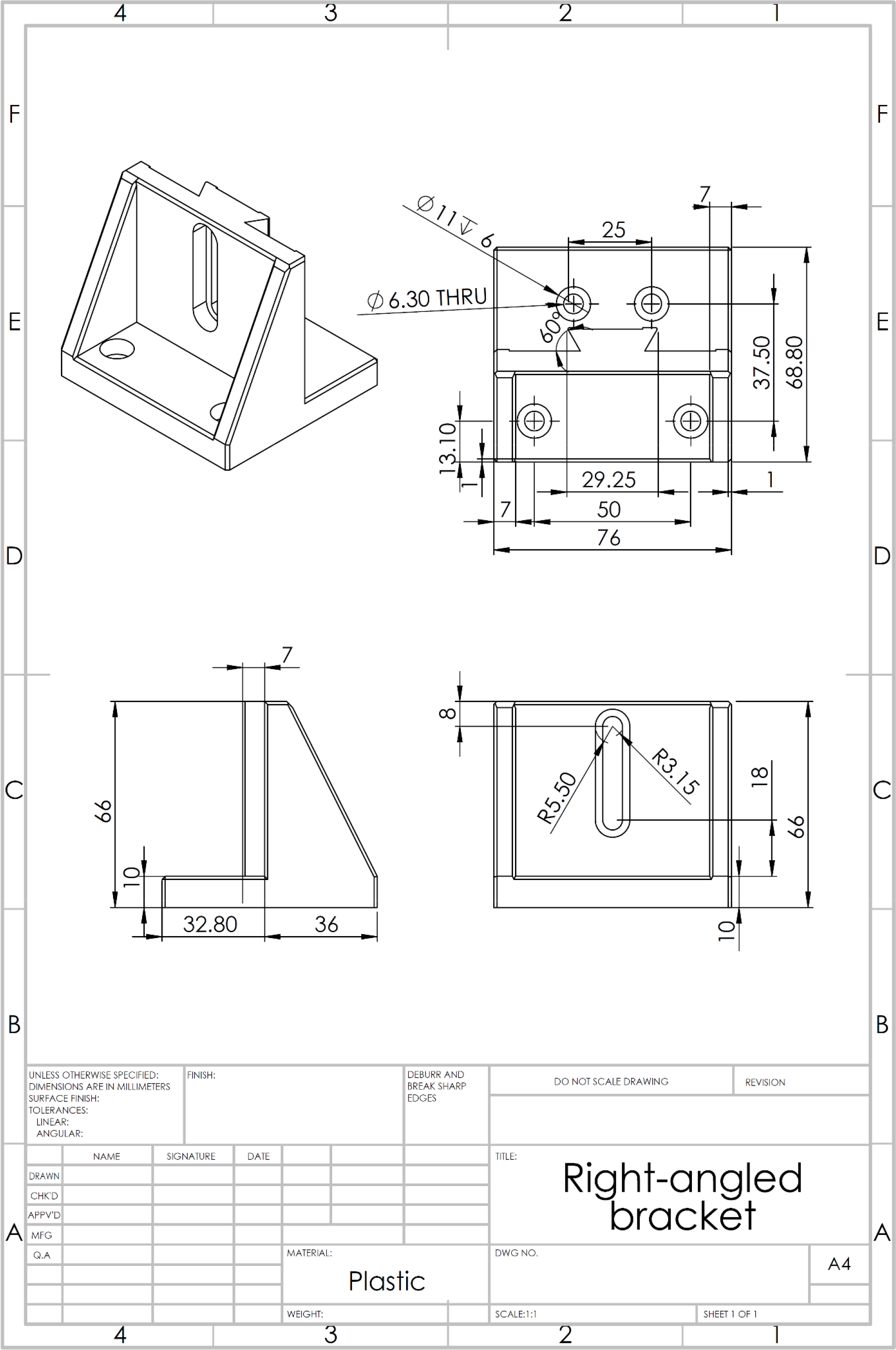

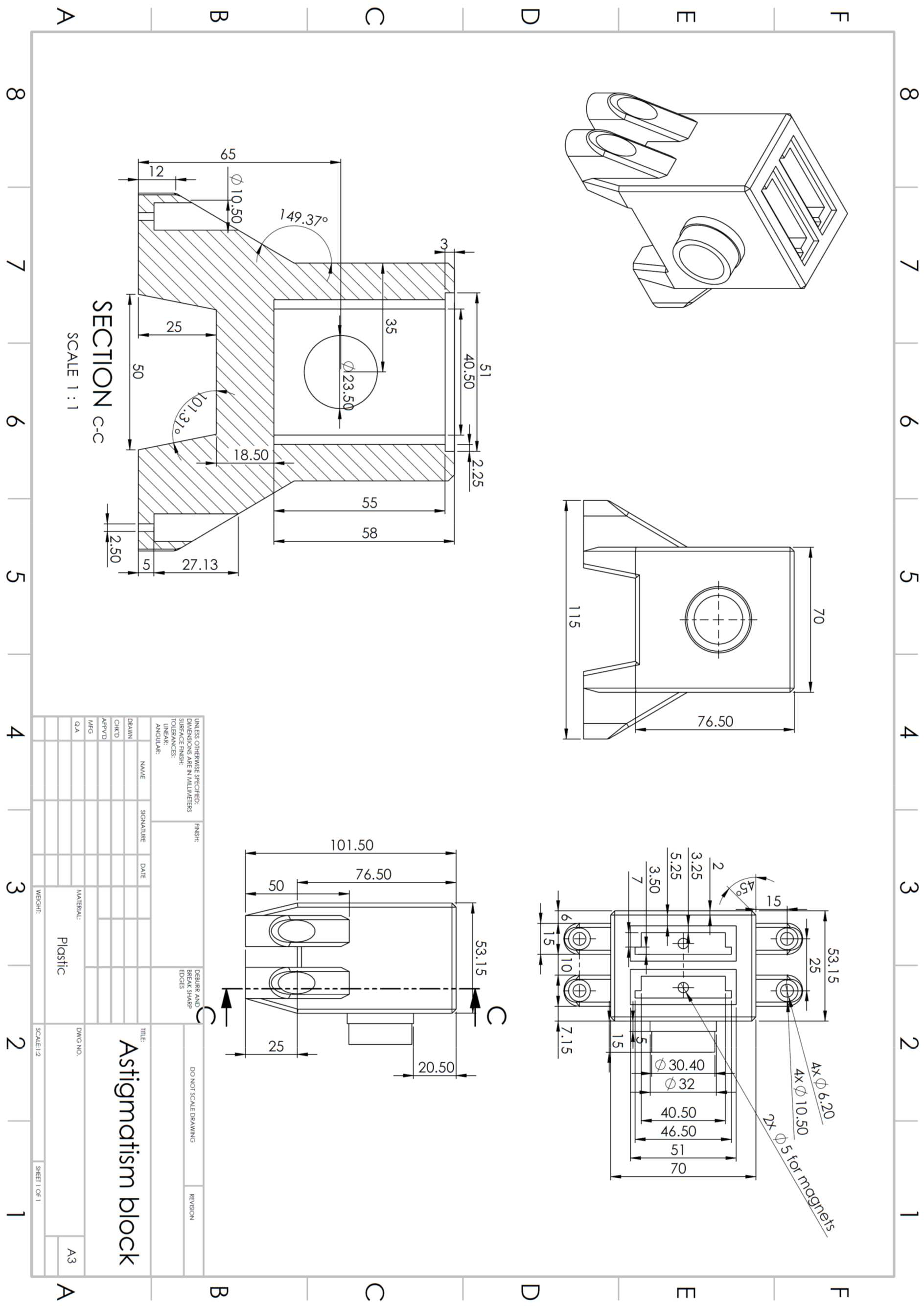

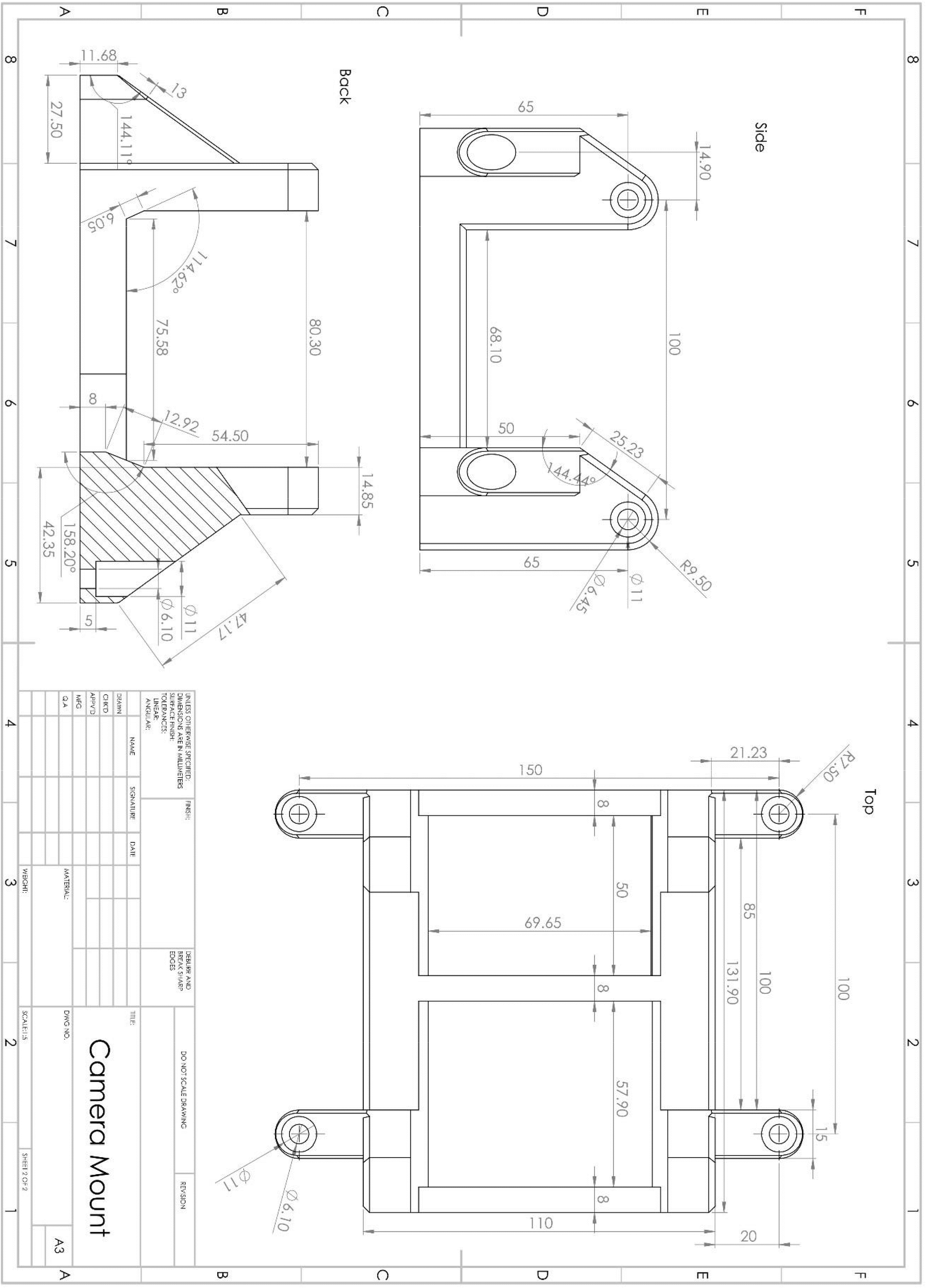

